# BAP1 loss and *PRAME* expression converge to remodel the tumor–immune ecosystem during uveal melanoma progression

**DOI:** 10.64898/2026.09.23.752454

**Authors:** James J. Dollar, Xiao Liu, Mithalesh Singh, Mahsa Sorouri, Jeffim N. Kuznetsoff, Sebastian Salazar, Anthony Cruz, Christina Decatur, Zelia M. Correa, J. William Harbour

## Abstract

Uveal melanoma (UM) is characterized by a small number of recurrent genetic alterations that determine metastatic propensity. BAP1 loss and *PRAME* expression define the dominant prognostic axes in UM, yet how they promote malignant progression remains unclear. We profiled 190,535 cells from normal uvea, uveal nevus, primary and metastatic UM using single-cell transcriptomics, T cell receptor sequencing, spatial transcriptomics and isogenic perturbation models. Normal melanocytes, nevus cells and UM cells formed a transcriptional continuum marked by loss of differentiation and emergence of neural crest-like, stress-responsive, hypoxic-glycolytic and immune-interacting states. BAP1 loss and *PRAME* expression imposed distinct but convergent immunoregulatory programs, inducing interferon and TNF–NF-κB signaling and MHC-I expression, with HLA-E showing the strongest response. These alterations were accompanied by macrophage and CD8^+^ T cell remodeling. PRAME-enriched tumor regions formed spatially organized niches enriched for macrophages and plasma cells. These findings define BAP1 loss and PRAME expression as distinct but convergent axes of tumor–immune coevolution and nominate HLA-E as a candidate mediator of immune resistance.

## INTRODUCTION

Cancer progression is increasingly understood to involve reciprocal coevolution between malignant cells and the immune microenvironment. While antitumor immunity can shape tumor evolution, how tumor genetic alterations in turn remodel the immune ecosystem remains poorly understood. Uveal melanoma (UM), a highly metastatic cancer of the eye, provides a highly tractable model for dissecting this process. Initiating G_αq_ pathway mutations arise early but are insufficient for malignant transformation, which requires additional, ordered molecular events. Among these, biallelic BAP1 loss drives the transition from the low-risk Class 1 to the high-risk Class 2 gene expression profile, whereas aberrant re-expression of the cancer–testis antigen PRAME increases metastatic risk independently of molecular class. Concurrently, UM presents an immunologic conundrum in which immune infiltration increases with malignant progression while failing to produce effective antitumor immunity, and metastatic disease is largely refractory to immune checkpoint blockade ^1^. Although the bispecific T cell engager tebentafusp improves survival in HLA-A*02:01-positive metastatic UM, the benefit is modest and unsustained in most patients^2^.

The immune resistance of UM is often attributed to its low tumor mutation burden^3^ and immune-privileged ocular environment^4^, yet neither explains why high-risk primary and metastatic tumors contain clonally expanded, melanoma-reactive T cells and many metastases harbor tumor-reactive tumor-infiltrating lymphocytes that nevertheless fail to mount an effective immune response^5–7^. We therefore hypothesized that BAP1 loss and *PRAME* expression contribute to immune resistance by reprogramming malignant and immune cell states. While BAP1 loss has been linked to inflammatory and immunosuppressive markers^8–10^, PRAME has largely been viewed as a prognostic biomarker and therapeutic target antigen. Whether PRAME actively reprograms malignant and immune cells, how its effects intersect with those of BAP1 loss, and whether these programs are spatially organized remain unknown.

Here, we addressed these questions by integrating single-cell transcriptomics, T cell receptor sequencing, spatial transcriptomics and isogenic perturbation models to reconstruct UM evolution using primary and metastatic UM samples, as well as normal uveal tissue and nevus samples. We show that BAP1 loss and *PRAME* expression exert distinct but convergent effects, with both activating interferon, TNF–NF-κB and MHC class I programs in tumor cells and HLA-E showing the most consistent induction among MHC class I genes. BAP1 loss and *PRAME* expression were also associated with distinct but complementary remodeling of macrophage and T cell states, while *PRAME*-expressing tumor cells were spatially associated with niches enriched for macrophages and plasma cells. Together, these findings establish a framework in which genetic and transcriptional changes associated with malignant evolution reshape the composition, functional state and spatial architecture of the immune microenvironment, and nominate HLA-E as a candidate mediator of immune resistance.

## RESULTS

### Single-cell transcriptional atlas of uveal melanoma

We generated a scRNA-seq atlas of 190,535 cells derived from 36 human samples, including 18 primary UM samples (122,560 cells), 8 metastatic UM samples (32,455 cells), 1 congenital uveal nevus (1,702 cells), 5 normal adult uveal samples (12,394 cells), and 4 normal fetal uveal samples (21,424 cells)(**Fig. 1a** and **Supplementary Table 1**). Unsupervised UMAP analysis revealed subtype-specific clustering of both neoplastic and nonneoplastic cells, segregating according to the 15-GEP/PRAME-defined subtype of the tumor from which they were derived^11^ (**Fig. 1b**). Lineage-specific marker genes were identified across all major cell populations (**Extended Data Fig. 1a** and **Supplementary Table 2**).

**Figure 1.**
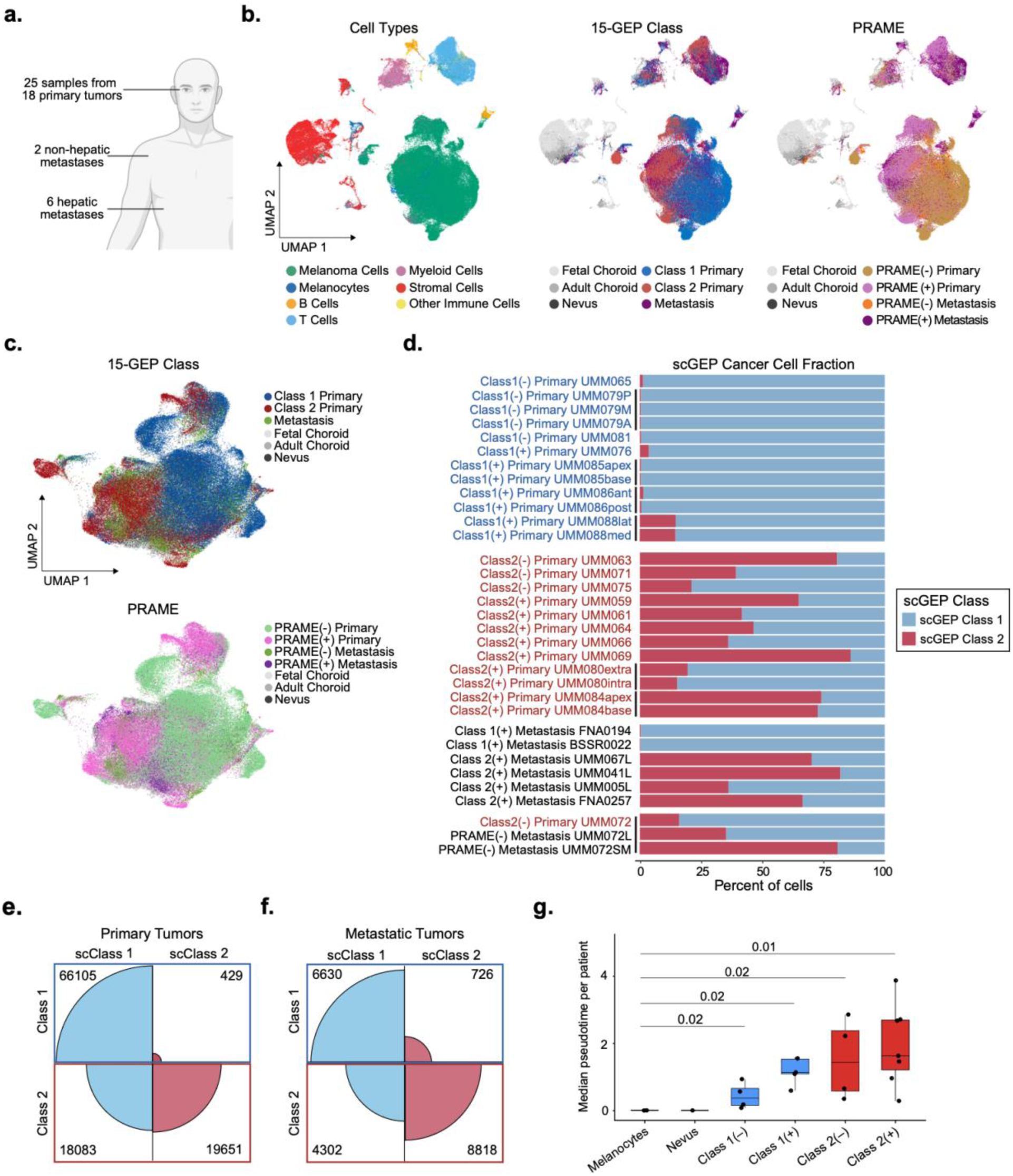
Single-cell transcriptional atlas of uveal melanoma. **a,** Schematic diagram of primary and metastatic uveal melanoma (UM) samples collected for single-cell RNA sequencing (scRNA-seq). **b,** Dimensionality reduction by Uniform Manifold Approximation and Projection (UMAP) plots of 190,535 cells annotated by major cell type, 15-gene expression profile (15-GEP) class, and *PRAME* expression status (negative or positive). **c**, UMAP plot of normal fetal (*n =* 1,427) and adult (*n =* 2,593) choroidal melanocytes and UM cells (*n =* 124,744) annotated by 15-GEP and *PRAME* status. **d,** Stacked bar plot depicting the relative abundance of support vector machine (SVM)-based inference of single-cell 15-GEP (scGEP) status for UM cells for each sample. Vertical black bars indicate samples originating from the same patient. Detailed summary of results available in Source Data. **e-f**, Confusion matrix plots displaying the concordance between UM cells based on 15-GEP clinical testing of the primary tumor (Class 1 or Class 2) versus the inferred scGEP (scClass1 or scClass2). Metastatic tumors were assigned the 15-GEP class of the patient-matched primary tumor. Blue and red quadrants represent inferred scClass 1 and scClass 2, respectively. **g**, Box-and-whisker plots showing median pseudotime per patient, grouped by molecular tumor subgroups, including adult choroidal melanocytes (*n =* 5), congenital nevus cells (*n =* 1), and Class 1/PRAME(–) (*n =* 4), Class 1/PRAME(+) (*n =* 5), Class 2/PRAME(–) (*n =* 4) and Class 2/PRAME(+) (*n =* 7) primary tumor cells. Boxes denote median and interquartile range (IQR), with whiskers extending to 1.5× IQR. Points represent individual patients’ median cellular pseudotime. Statistical significance was assessed using two-sided Wilcoxon rank-sum test. Benjamini-Hochberg (BH)-adjusted *P* values are shown for inter-group comparisons.

### Single-cell landscape of normal and neoplastic human uveal melanocytes

We then analyzed normal and neoplastic melanocytes across all 36 samples, including 124,744 UM cells, 204 nevus cells, 2,389 normal adult uveal melanocytes (UMCs) and 1,427 normal fetal UMCs (**Fig. 1c**). As anticipated, unsupervised hierarchical clustering of pseudobulk transcriptomes from 18 primary UMs (including five sampled across multiple regions) revealed segregation by 15-GEP class, with multiregional specimens from the same tumor clustering closely together (**Extended Data Fig. 1b**). We trained an SVM algorithm to infer the 15-GEP Class (scGEP) of single tumor cells as Class 1 (scClass1) or Class 2 (scClass2) based on expression of the 12 differentially expressed genes (DEGs) from the 15-GEP (**Extended Data Fig. 1c**). Although this approach cannot fully replicate the clinical 15-GEP assay, which uses machine learning and a training set to analyze genes that are expressed in both tumor and immune cells^5,12^, it allowed us to estimate the proportion of scClass1 and scClass2 UM cells in each tumor sample as a means of investigating tumor heterogeneity and evolution. Whereas Class 1 tumors were comprised almost entirely of scClass1 cells, Class 2 tumors contained nearly equal proportions of scClass1 (49.9%) and scClass2 (50.1%) cells (**Fig. 1d-f**). Metastatic tumor samples arising from two different Class 1 primary tumors were comprised almost exclusively of scClass1 tumor cells (92.3%), whereas metastatic tumor samples arising from six different Class 2 primary tumors contained a mixture of scClass1 (39.3%) and scClass2 (60.7%) cells. In one case (UMM072), a Class 2 primary tumor and two metastatic samples were available for analysis and showed an enrichment for scClass2 cells during progression from the primary tumor to an initial liver metastasis and to a later subcutaneous metastasis. Taken together, these findings are consistent with previous work showing low intratumoral heterogeneity for the 15-GEP within primary tumors^12^, and they support a model in which Class 2 tumors arise from Class 1 tumors, with BAP1 loss conferring a selective advantage that drives clonal expansion and eventual dominance of BAP1-deficient tumor cells.

### Recurrent transcriptional states in uveal melanoma

We applied Monocle3 pseudotime analysis to determine whether normal and neoplastic uveal melanocytes could be ordered along a putative transcriptional continuum, with normal adult uveal melanocytes designated as the root population. Consistent with this hypothesis, median pseudotime values were comparable between normal uveal melanocytes and uveal nevus cells but increased significantly in a stepwise manner across subgroups from Class 1/PRAME(–) to Class 1/PRAME(+), Class 2/PRAME(–), and Class 2/PRAME(+) tumors (**Fig. 1g**).

Using a cluster-based, marker-guided approach, we found that UM cells segregated into recurrent transcriptional states which differ from the canonical MITF-low/AXL-high framework described in cutaneous melanoma^13^. These states include a baseline differentiated state, as well as proliferative, hypoxic-glycolytic, vasculogenic, neural crest-like, stress-responsive and immune-interacting states (**Fig. 2a-d**). The differentiated state was the most abundant overall and was enriched in low-risk Class 1/PRAME(–) primary tumors, with progressively lower abundance across PRAME(+), Class 2, and metastatic tumors (**Extended Data Fig. 2**).

**Figure 2.**
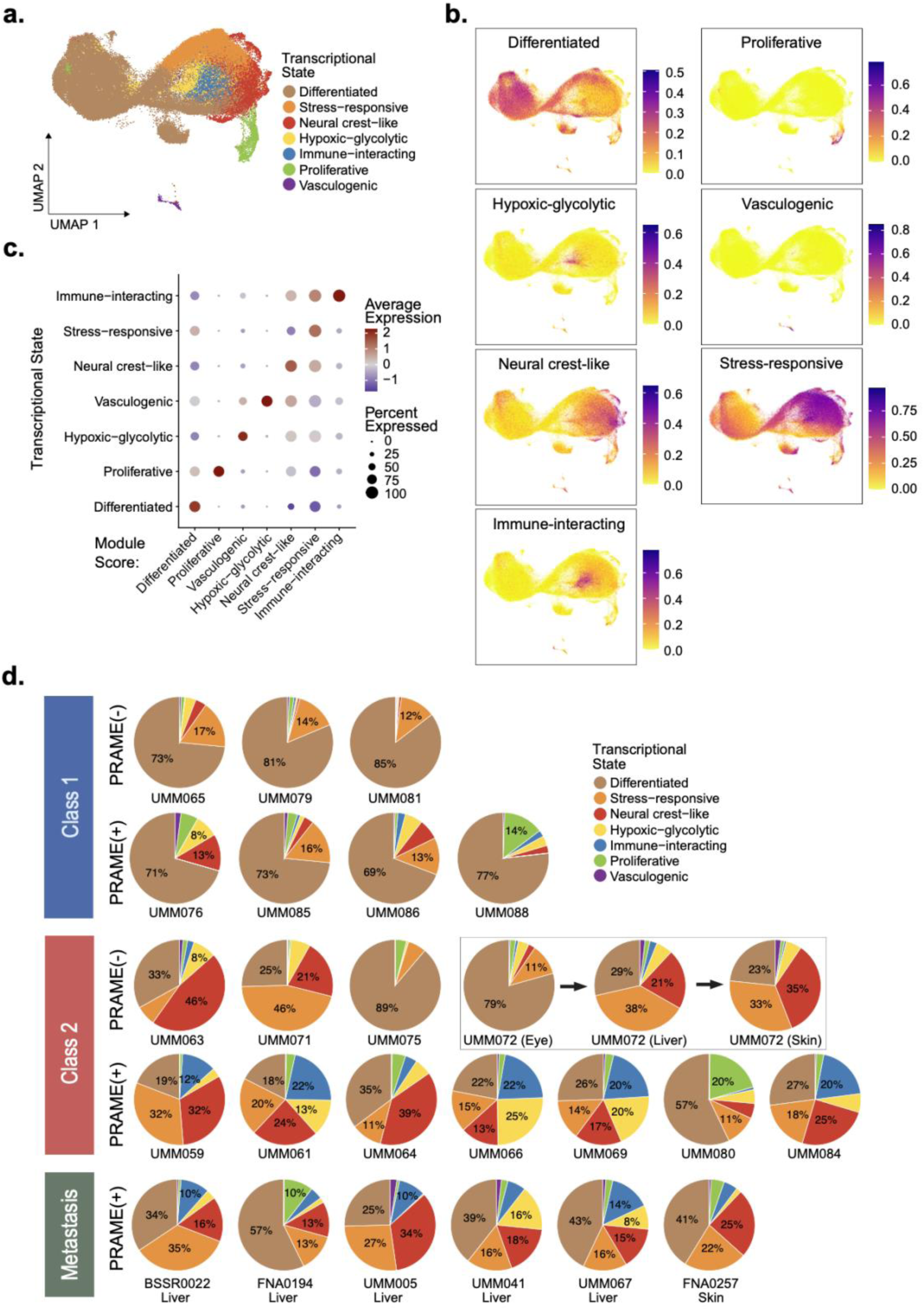
Transcriptional states in primary and metastatic uveal melanoma. **a,** Uniform Manifold Approximation and Projection (UMAP) plot of 122,715 strictly filtered tumor cells from 25 primary and 8 metastatic uveal melanoma (UM) samples, with each cell colored according to its cluster-defined transcriptional state: differentiated, hypoxic-glycolytic, neural crest-like, immune-interacting, proliferative, vasculogenic or stress-responsive. **b,** Distribution of the seven transcriptional state module scores on the same UMAP plot, calculated with UCell using curated gene sets (**Supplementary Table 3**). Color scales were determined independently for each module. **c**, Dot plot of module scores (x-axis) across discrete transcriptional states (y-axis). Dot size indicates the percent of cells expressing the gene program; dot color indicates the average scaled expression. **d,** Tumor-level summary pie charts depicting the transcriptional state composition of primary UM samples grouped by 15-GEP class and *PRAME* expression status, and metastatic UM samples grouped by *PRAME* status. Multiregional samples from the same tumor were combined for UMM079, UMM080, UMM084, UMM085, UMM086 and UMM088. Box highlights matched primary, liver and subcutaneous metastatic tumors for UMM072.

Reciprocally, the neural crest-like state was virtually absent in Class 1/PRAME(–) tumors and became progressively enriched across advancing molecular classes, reflecting a shift towards less differentiated states with tumor evolution. The hypoxic-glycolytic and stress-responsive states also exhibited progressive enrichment with malignant evolution, whereas the immune-interacting state was largely confined to Class 2/PRAME(+) and metastatic tumors. In contrast, proliferative and vasculogenic states represented a small proportion of tumor cells and showed no consistent association with molecular subgroup or metastatic status. Metastatic UM cells recapitulated the same transcriptional states observed in primary tumors, with stress-responsive and neural crest-like states being most abundant and with no metastasis-specific states identified. InferCNV identified canonical UM chromosomal copy number variations (CNVs)^14,15^ in cells across all states (**Supplementary Fig. 1**), confirming that these were indeed malignant UM cells rather than contaminating normal cells. These findings support a model in which uveal melanoma evolves along a transcriptional trajectory away from a differentiated melanocytic state, accompanied by selective enrichment of a limited repertoire of recurrent transcriptional states associated with increasingly aggressive molecular subgroups and metastatic risk.

### BAP1 loss and *PRAME* expression drive distinct and convergent transcriptional reprogramming of uveal melanoma cells

Given the disproportionate contribution of BAP1 loss and aberrant *PRAME* expression to UM prognosis^11,16^, we analyzed single-cell transcriptomic signatures associated with each of these aberrations in UM cells. First, we identified DEGs in UM cells from Class 2 versus Class 1 primary tumors, irrespective of *PRAME* status, and performed gene set enrichment analysis (GSEA)(**Fig. 3a-b**). UM cells from Class 2 tumors exhibited increased expression of established markers of high-risk UM (e.g., *HTR2B, ECM1, PROS1*)^8,12^, together with activation of transcriptional programs associated with cellular stress adaptation (e.g., *GDF15, CHAC1*), extracellular matrix remodeling and invasion (e.g., *PTP4A3, HTRA1*), vascular morphogenesis (e.g., *CCN1, DLL4*), TNF/NF-κB signaling (e.g., *NFKBIA, IL1R1*), interferon response (e.g., *ISG15, WARS*), and MHC class I antigen presentation (e.g., *HLA-B, TAP1*), while programs involved in protein translation, ribosome biogenesis, and RNA processing were concomitantly attenuated. Interestingly, genes upregulated in Class 2 tumors included *CYSLTR2* and *PLCB4*, which are recurrently mutated in UM and function as activators of oncogenic G_q/11_ signaling^17,18^, raising the possibility of crosstalk between G_q/11_ and BAP1 pathways. We next identified DEGs and performed GSEA in UM cells from PRAME(+) versus PRAME(–) tumors irrespective of 15-GEP class (**Fig. 3c–d**). PRAME(+) UM cells exhibited upregulation of GDF15, as well as genes involved in interferon and NF-κB signaling, MHC class I and II antigen presentation, and other immune regulatory pathways.

**Figure 3.**
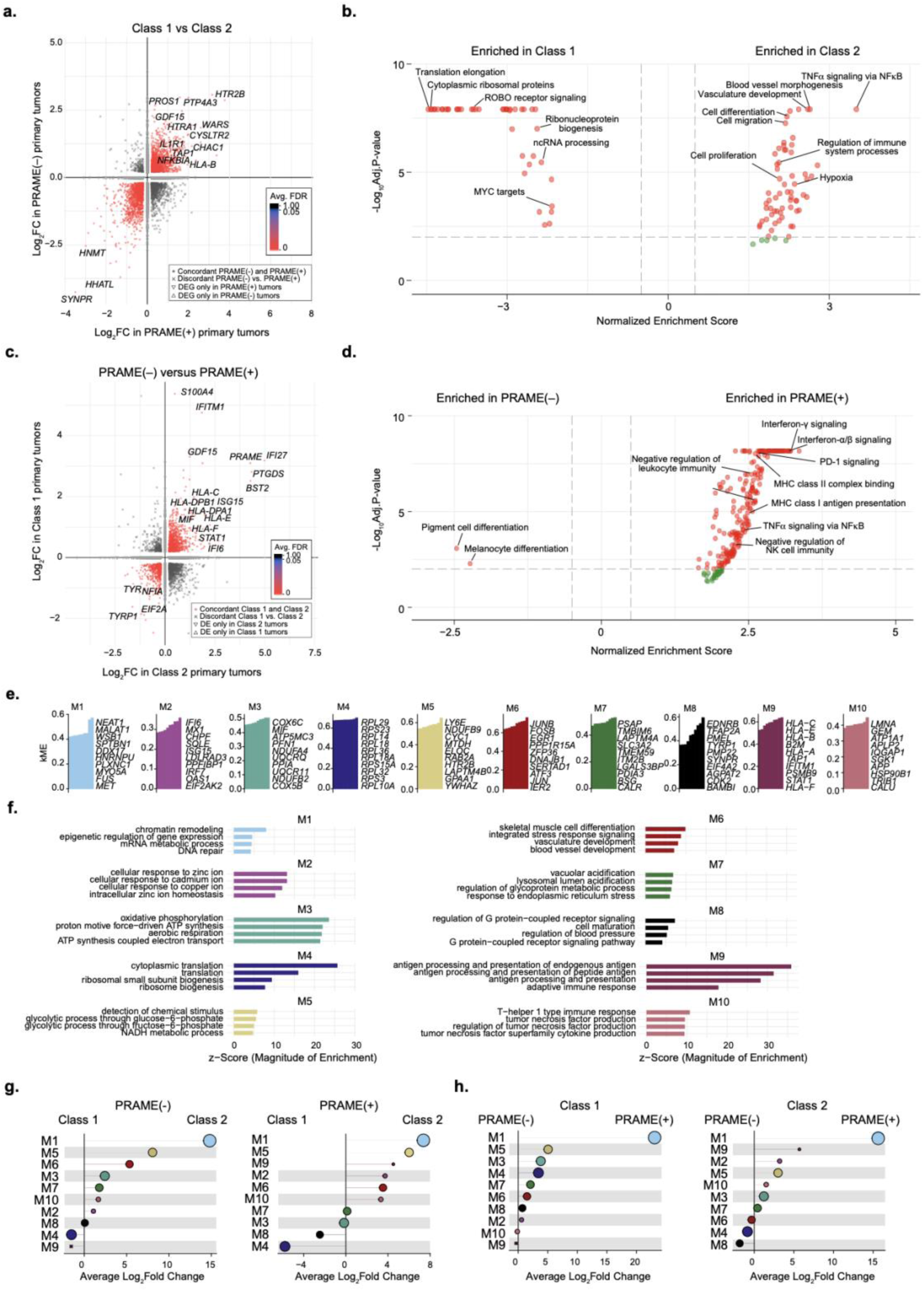
Shared and distinct transcriptional programs associated with 15-GEP class and *PRAME* expression status in uveal melanoma cells. **a,** Double-volcano plot depicting differentially expressed genes (DEGs) in uveal melanoma (UM) cells from Class 1 (*n =* 7) versus Class 2 (*n =* 11) primary tumors irrespective of *PRAME* status (FDR < 0.05 and absolute log_2_FC > 0.2). Genes upregulated in Class 2 relative to Class 1 tumors in both PRAME(+) and PRAME(–) tumors are indicated in the upper right quadrant, whereas those downregulated in Class 2 relative to Class 1 tumors in both PRAME(+) and PRAME(–) tumors are indicated in the lower left quadrant. Key genes mutually upregulated (upper right quadrant) or downregulated (lower left quadrant) across both axes are highlighted. Detailed DEG analysis results are available in Source Data. **b**, Volcano plot showing gene set enrichment analysis (GSEA) results for DEGs in Class 2 versus Class 1 tumors, irrespective of *PRAME* status. Detailed GSEA results are available in Source Data. **c**, Double-volcano plot depicting DEGs in PRAME(+) (*n =* 11) versus PRAME(–) primary tumors (*n =* 7) irrespective of 15-GEP class status. Genes upregulated in PRAME(+) relative to PRAME(–) tumors in both Class 1 and Class 2 tumors are indicated in the upper right quadrant, whereas those downregulated in PRAME(+) relative to PRAME(–) tumors in both Class 1 and Class 2 tumors are indicated in the lower left quadrant. Key genes mutually upregulated (upper right quadrant) or downregulated (lower left quadrant) across both axes are highlighted. **d**, Volcano plot showing GSEA results for DEGs in PRAME(+) versus PRAME(–) primary tumors, irrespective of Class status. Detailed GSEA results are available in Source Data. **e**, Weighted gene co-expression network analysis (WGCNA) of 128,764 normal and neoplastic melanocytes, comprising 2,389 adult choroidal melanocytes (5 patients, 13 samples), 1,427 fetal choroidal melanocytes (4 patients, 8 samples), 204 congenital uveal nevus cells (1 sample), and 124,744 UM cells from primary tumors (104,268 cells; 25 samples; 18 patients) and metastatic tumors (20,476 cells; 8 samples; 7 patients). The top 10 melanocytic co-expression gene modules (M1–M10) are shown, with their top hub genes plotted as vertical bars along the x-axis and listed to the right of each plot. The y-axis represents the intramodular eigengene-based connectivity score (kME) for each hub gene. **f**, Bar plots depict the top enriched gene ontology biological processes for each module (*P* value < 0.05). The x-axes display z -scores representing the magnitude of enrichment for each biological process. **g**, Differential module score analysis for Class 2 versus Class 1 primary tumors separated by *PRAME* expression status. **h**, Differential module score analysis for PRAME(+) versus PRAME(–) primary tumors separated by 15-GEP class. Module scores were generated with UCell using the first 10 hub genes and two-tailed Wilcoxon test.

Conversely, genes downregulated in PRAME(+) tumors were enriched for melanocyte differentiation programs, consistent with the stem-like dedifferentiated states described above. Together, these findings suggest that BAP1 loss and *PRAME* expression represent complementary axes driving transcriptional evolution in UM toward a tumor cell-intrinsic immunosuppressive state, with both genetic aberrations associated with immunomodulatory and inflammatory signaling, BAP1 loss further promoting chronic stress adaptation and vascular remodeling, and *PRAME* expression enhancing inflammatory signaling, antigen presentation, and expression of immunomodulatory mediators.

Using Weighted Gene Co-expression Network Analysis (WGCNA), an orthogonal unsupervised approach, we identified 10 core transcriptional modules operating within the UM cell population (**Fig. 3e-h**). A chromatin/RNA-regulatory module (M1) was the most consistently enriched module in both Class 2 and PRAME(+) tumors, indicating a shared transcriptional remodeling program in high-risk UM. Class 2 tumors were also enriched for metabolic (M3, M5), stress-response (M6, M7), and immune-related (M9, M10) modules. Of note, the antigen-presentation module (M9), which was most enriched in Class 2/PRAME(+) tumors, was particularly marked by an increase in expression of MHC class I genes, especially the nonclassical class I molecule *HLA-E.* These findings indicate that the transcriptional consequences of BAP1 loss and *PRAME* expression reflect coordinated regulatory reprogramming rather than isolated gene-level alterations and suggest that phenotypic plasticity, stress adaptation, and immune modulation are key drivers of metastatic progression in UM.

To determine whether the transcriptional programs associated with Class 2 and PRAME(+) tumors might be directly driven by BAP1 loss and *PRAME* expression, respectively, we experimentally modulated BAP1 and *PRAME* levels in engineered UMCs and UM cell lines (**Extended Data Fig. 3a-b** and **Supplementary Fig. 2**). Following CRISPR-mediated BAP1 knockout or doxycycline-inducible BAP1 knockdown in three independent isogenic models, BAP1 depletion consistently induced interferon-stimulated genes (*ISG15, IFIT1* and *IFI27*), TNF/NF-κB signaling components (*TNFA, NFKB1, NFKB2, RELA* and *RELB*), and MHC class I genes (*HLA-A, HLA-B* and *HLA-E*) (**Extended Data Fig. 3a**). Although induction of *IFNB* varied between models, downstream canonical interferon-responsive genes were consistently induced. Among MHC class I genes, *HLA-E* exhibited the strongest and most consistent induction across all three models, substantially exceeding the more modest changes observed for the classical class I genes *HLA-A* and *HLA-B*, while *HLA-C* remained unchanged or was slightly reduced. We next evaluated the same panel of genes following *PRAME* induction in engineered UMCs and UM cells (**Extended Data Fig. 3b**). Compared with BAP1 loss, *PRAME* induction resulted in more consistent upregulation of *IFNB*, robust induction of canonical interferon-stimulated genes, activation of the TNF/NF-κB axis, and consistent upregulation of all four MHC class I genes, with *HLA-E* remaining the most strongly induced. Together, these findings confirm that BAP1 loss and *PRAME* expression converge on overlapping interferon, TNF/NF-κB and MHC-I antigen-presentation programs, and they indicate that *PRAME* expression results in broader and more consistent activation of these pathways. Notably, *HLA-E* was among the most strongly induced MHC class I genes following both BAP1 depletion and *PRAME* induction, supporting a potential role as a mediator of tumor–immune interactions in high-risk UM.

### BAP1 loss and *PRAME* expression drive extensive remodeling of the tumor immune microenvironment

Given the extensive transcriptional reprogramming associated with BAP1 loss and *PRAME* expression, we next investigated their effects on the tumor immune microenvironment (TIM). Overall, the abundance of immune cells progressively increased from Class 1 to Class 2 primary tumors, PRAME(–) to PRAME(+) primary tumors, and primary to metastatic tumors (**Fig. 4a**). To identify local cell neighborhoods enriched in Class 2 and PRAME(+) tumors, we performed differential abundance analysis using MiloR^19^. Class 2 and PRAME(+) tumors were both enriched CD8+ tissue-resident memory T cells (CD8.TRM), CD8+ effector memory T cells (CD8.EM) and exhausted CD8+ T cells (CD8.TEX)(**Fig. 4b**). Class 2 tumors were enriched for SPP1^+^ macrophages, and PRAME(+) tumors were enriched for CD4^+^ naïve T cells, NK cells and CD4^+^ regulatory T cells (CD4.TREG). Class 1/PRAME(+) tumors were specifically enriched for B lineage cells.

**Figure 4.**
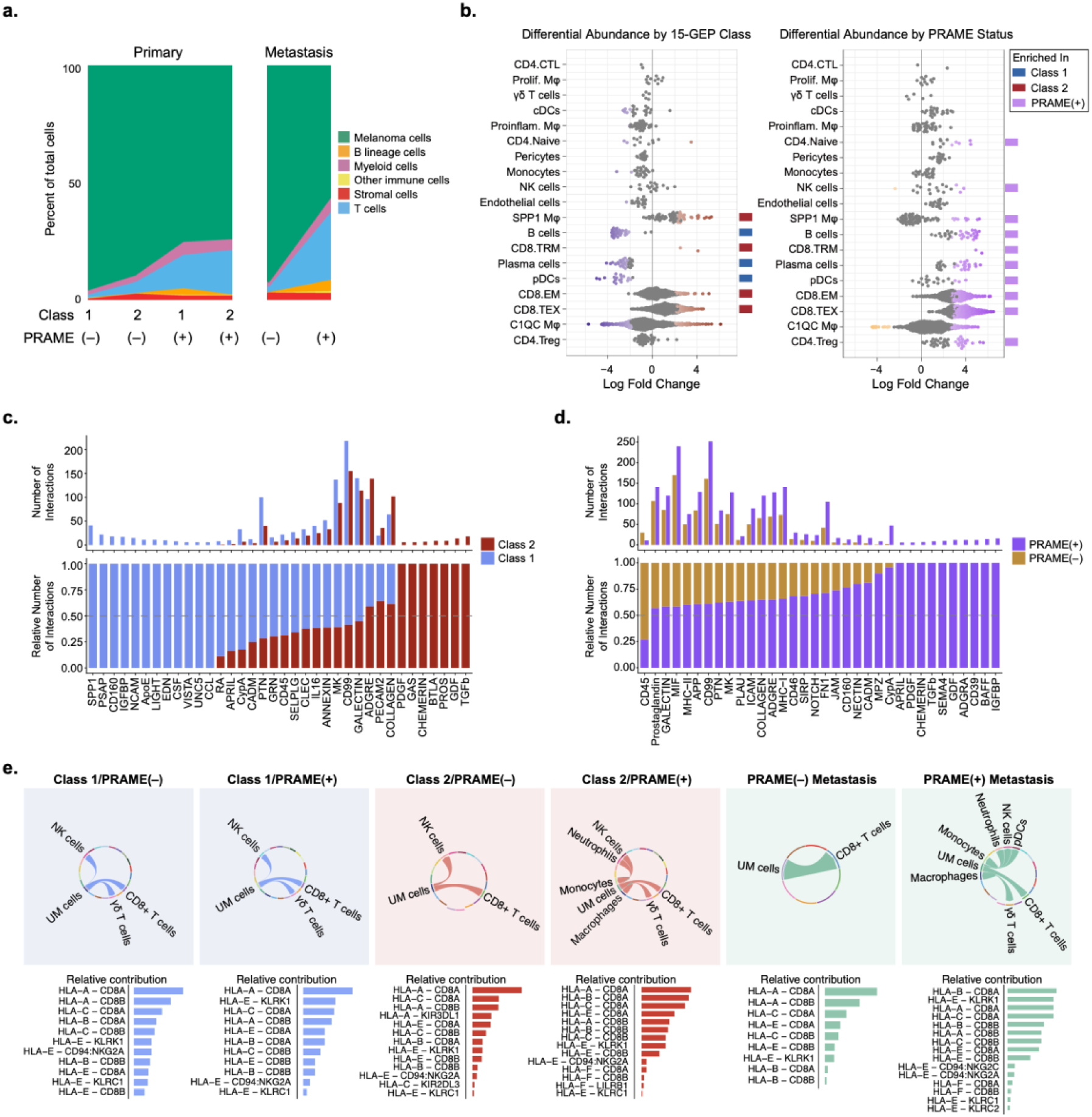
Immune cell composition of the tumor microenvironment across uveal melanoma molecular subgroups. **a**, Stacked-area plot exhibiting relative abundance of the major immune cell types by 15-GEP class for primary tumors and *PRAME* status for primary and metastatic tumors. **b**, Swarm plot of differentially abundant neighborhoods (FDR < 0.05) by cell type based on 15-GEP class and *PRAME* status. Differential abundance of cellular neighborhoods was determined by negative binomial generalized linear modeling with MiloR. Blue, red and purple boxes indicate enrichment in Class 1, Class 2 and PRAME(+) tumors, respectively. **c-d**, Ranked net plots of significant overall signaling networks within the tumor immune microenvironment, comparing ligand-receptor interactions in **c**, Class 2 versus Class 1 primary tumors and in **d**, PRAME(–) versus PRAME(+) primary tumors. Signaling networks were considered significant at *P* < 0.01 and a scaled contribution > 5 in at least one compared group. Full signaling networks for tumor groups are available in Source Data. The upper bar plots show the absolute number of ligand-receptor interactions, while the lower stacked bars display the relative number of ligand-receptor interactions, highlighting the proportional contribution of each pathway to the overall communication landscape in Class 1 (blue) and Class 2 (red) tumors, and in PRAME(–) (orange) and PRAME(+) (purple) tumors. **e**, Chord plots depicting the CellChat-predicted MHC-I signaling network originating from UM cells, with chords linking UM cells to inferred target cell types. The corresponding bar plot beneath each chord plot summarizes the predicted ligand-receptor interactions ranked by relative contribution for each molecular tumor subgroup.

Using CellChat to infer ligand-receptor and cell-cell communications within the TIM, we found that both BAP1 loss and *PRAME* expression were associated with extensive remodeling of intercellular signaling networks (**Fig. 4c-d**). Rather than altering a single dominant pathway, these alterations were accompanied by coordinated shifts across multiple signaling programs, indicating progressive reorganization of tumor-immune communications. Transition from Class 1 to Class 2 tumors was characterized by a shift from pleiotrophin (PTN)- to prostaglandin-dominant signaling, consistent with a more inflammatory and immune-regulatory microenvironment^20–22^. Similarly, PRAME(–) tumors preferentially utilized CXCL-, CD45-, and CLEC-mediated signaling, consistent with antitumor immune recognition and surveillance^23–25^, whereas PRAME(+) tumors exhibited increased ADGRE-family signaling, which has been linked to TIM dysfunction^26^. These findings suggest that molecular progression is accompanied not only by remodeling of tumor cell transcriptional states and immune cell composition, but also by progressive reorganization of intercellular communications within the TIM.

Considering the increased expression of HLA class I genes in PRAME(+) UM cells (**Fig. 2c–d**) and the strong induction of HLA class I genes following ectopic PRAME expression in experimental models (**Extended Data Fig. 3b**), we next investigated whether these transcriptional changes were accompanied by remodeling of HLA-mediated tumor–immune communication (**Fig. 4e**). HLA-A/B/C–CD8A/CD8B interactions predominated across all molecular subgroups, consistent with preservation of antigen presentation to cytotoxic T cells throughout malignant evolution. However, the HLA communication network became progressively more complex with increasing molecular risk. Class 1 tumors exhibited HLA signaling primarily between UM cells and cytotoxic lymphocytes, whereas Class 2 tumors incorporated additional interactions with NK cells and myeloid populations. Notably, HLA-E-mediated interactions became progressively more prominent with malignant evolution, particularly those involving the inhibitory NK-cell receptor complex CD94/NKG2 (KLRD1/KLRC1). PRAME(+) metastatic UM cells exhibited the most diverse HLA communication network, incorporating interactions with macrophages, monocytes, neutrophils, NK cells and T cells. Collectively, these findings suggest that BAP1 loss and *PRAME* expression are associated not only with global remodeling of tumor–immune communication but also with progressive diversification of HLA-mediated signaling, highlighting HLA-E as a candidate mediator of immune regulation during uveal melanoma progression.

### BAP1 loss and *PRAME* expression cooperatively remodel macrophage transcriptional states

We then examined the myeloid compartment, where cells clustered according to cell type, including monocytes, macrophages, and other myeloid cell types (**Fig. 5a**). Within each cluster, cells tended to stratify according to 15-GEP class and *PRAME* status. Given the predominance of macrophages within the myeloid compartment, we focused subsequent analyses on this lineage. DEG analysis identified genes differentially expressed in tumor-associated macrophages (TAMs) between Class 1 and Class 2 tumors irrespective of *PRAME* status, as well as those between PRAME(–) and PRAME(+) tumors, irrespective of 15-GEP class (**Fig. 5b**). TAMs from Class 1 tumors preferentially expressed genes associated with an active innate immune response (e.g., *IFIT1, IFITM1, CX3CR1, CD37, C3*), whereas Class 2 TAMs were enriched for immune-regulatory genes (e.g., *CXCL1, CCL20, IL6, LAG3, CD209*). TAMs from PRAME(–) tumors preferentially expressed genes associated with a chemokine-rich inflammatory program (e.g., *IL1A, CXCL1, CXCL3, CXCL8, CCL20*), whereas those from PRAME(+) tumors were characterized by expression of canonical interferon-stimulated genes (e.g., *STAT1, IFI27, IFIT1, IFITM1, ISG15*), together with MHC class II genes (e.g., *HLA-DOB, HLA-DQA2*), suggesting broad transcriptional reprogramming toward an interferon-responsive, antigen-presenting phenotype. GSEA corroborated these findings, demonstrating enrichment of TNFα signaling via NF-κB and regulation of T cell activation in Class 2 TAMs and enrichment of type I and type II interferon signaling pathways in PRAME(+) TAMs (**Fig. 5c**). Collectively, these findings suggest that Class 2/BAP1-deficiency and *PRAME* expression independently shape orthogonal dimensions of macrophage transcriptional states that may cooperatively promote an immunosuppressive TIM.

**Figure 5.**
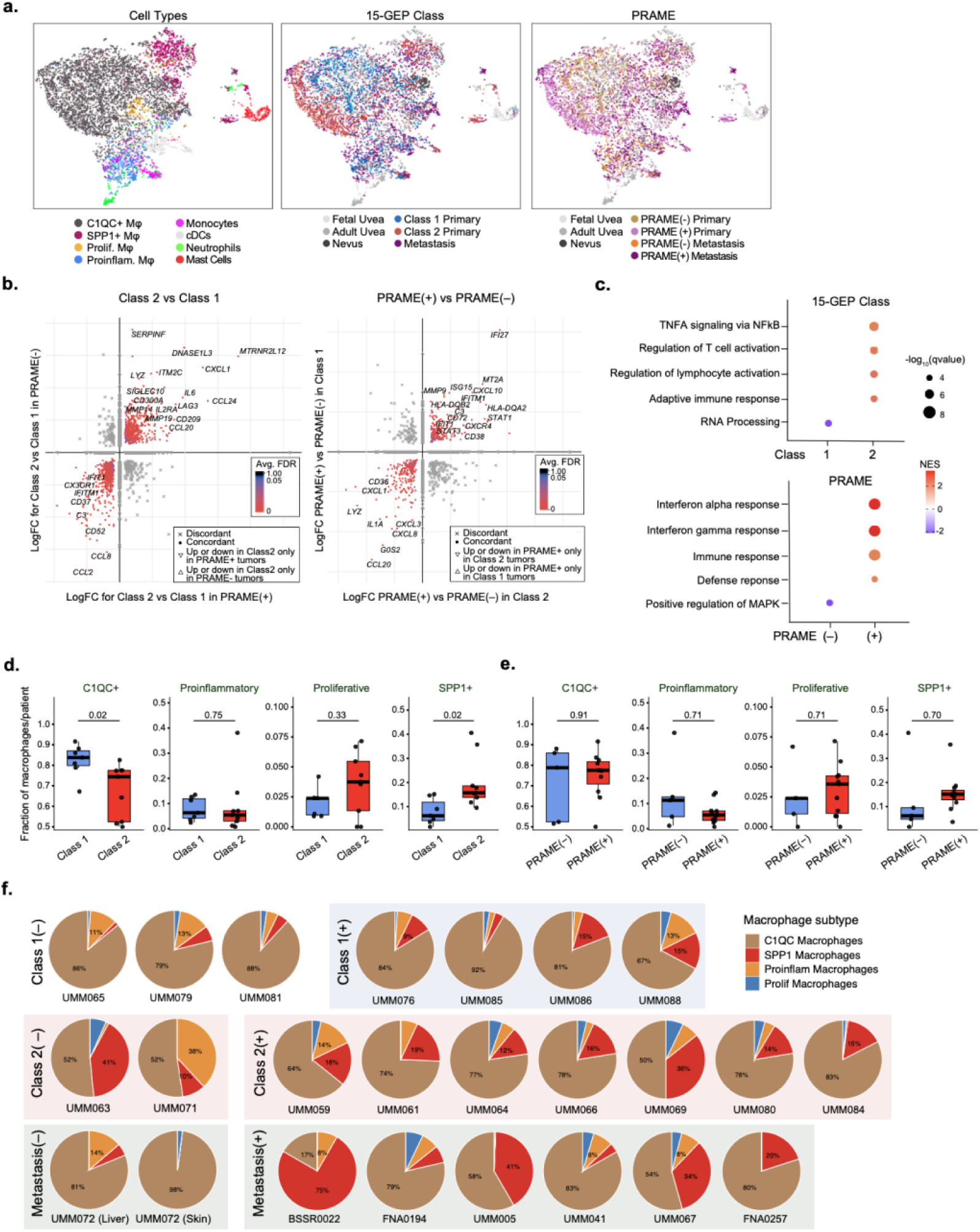
Macrophage transcriptional programs and subtype composition across uveal melanoma molecular subgroups. **a**, Uniform Manifold Approximation and Projection (UMAP) plots depicting myeloid cells (*n =* 6,979) annotated according to cell type, 15-GEP Class and *PRAME* expression status. **b**, Double-volcano plot showing log2 fold-change (log2FC) of significantly differentially expressed genes (DEGs) in tumor associated macrophages (TAM) from Class 2 versus Class 1 (left) and PRAME(+) versus PRAME(–) (right) primary tumors (FDR-adjusted *P* < 0.05, absolute log2FC > 0.2). In the left volcano plot, the x-axis shows log2FC for the Class 2 versus Class 1 comparison among PRAME(+) tumors, and the y-axis shows log2FC for the same comparison among PRAME(–) tumors. In the right volcano plot, the x-axis and y-axis show log2FC for the PRAME(+) versus PRAME(–) comparison within Class 1 and Class 2 tumors, respectively. Key genes mutually upregulated (upper right quadrant) or downregulated (lower left quadrant) across both axes are highlighted. Complete DEG analysis results available in Source Data. **c**, Dot plot showing the top hallmark gene set enrichment analysis (GSEA) results for DEGs in TAMs with respect to 15-GEP Class and *PRAME* expression status. Dot size indicates inverse log_10_ of q values, and color indicates normalized enrichment score (NES). **d,e**, Box-and-whisker plots comparing the relative abundance of each indicated major TAM subtype, expressed as a fraction of all macrophages per patient in **d**, Class 2 (*n* = 9) versus Class 1 (*n* = 7) and **e**, PRAME(+) (*n* = 11) versus PRAME(–) (*n* = 5) primary tumors. Boxes represent median and interquartile range (IQR), whiskers extend to 1.5× IQR, and each point represents one patient. Statistical significance was assessed using two-sided Wilcoxon rank-sum test and Benjamini-Hochberg (BH)-adjusted *P* values. **f**, Pie charts depicting the composition of TAM subtypes for each tumor grouped by 15-GEP class and *PRAME* expression status for primary tumors and *PRAME* status for metastatic tumors.

Previous studies have suggested that high-risk UMs are enriched for alternatively activated M2-like TAMs based on the traditional M1/M2 polarization framework^27^. However, neither M1 nor M2 macrophage polarization signatures were significantly associated with 15-GEP class or *PRAME* status in primary or metastatic tumors (**Supplementary Fig. 3**), indicating that the conventional *in vitro*-derived M1/M2 paradigm^28^ does not adequately capture TAM heterogeneity by scRNA-seq. Therefore, we evaluated macrophage transcriptional states using an scRNA-seq-based macrophage annotation framework^29^(**Fig. 5a, d-f**). To ensure comparison within a consistent tissue context, we restricted this analysis to primary tumors, thereby avoiding potential confounding by organ-specific microenvironments at metastatic sites. C1QC+ TAMs represented the dominant macrophage population across all primary tumors and showed a significantly reduced abundance in Class 2 compared to Class 1 tumors. In contrast, SPP1^+^ TAMs were significantly enriched in Class 2 primary tumors, consistent with our differential abundance analysis (**Fig. 4b**). By contrast, *PRAME* expression status was not significantly associated with any TAM subgroup (**Fig. 5e**), although there as a significant enrichment of SPP1^+^ TAMs in PRAME(+) tumors using neighborhood abundance analysis (**Fig. 4b**). Further, there was a trend towards increased abundance of SPP1^+^ TAMs with *PRAME* expression across metastatic tumors (**Fig. 5f**). These data indicate that SPP1 marks a biologically relevant TAM subset associated with high-risk UM.

### BAP1 loss and *PRAME* expression are associated with distinct T cell phenotypes and clonal architecture

Given the extensive remodeling of tumor cell and macrophage compartments associated with BAP1 loss (Class 2) and *PRAME* expression, we next examined whether these alterations were accompanied by corresponding changes in the adaptive immune compartment, focusing on T cell transcriptional states. Unsupervised UMAP clustering of tumor-infiltrating T cells identified transcriptionally distinct CD4^+^ and CD8^+^ T cell populations (**Fig. 6a**). CD4^+^ T cells comprised naïve, cytotoxic (CTL), T follicular helper (Tfh), and regulatory T cell (Treg) subsets, whereas CD8^+^ T cells resolved into central memory (CM), effector memory (EM), tissue-resident memory (TRM), terminal effector (TEMRA), and exhausted (CD8-TEX) populations. Discrete MAIT and γδ T-cell populations were also identified. UMAP visualization demonstrated a continuum of CD8^+^ T cell differentiation states extending from memory through effector to a large, exhausted T cell compartment, whereas CD4^+^ subsets formed transcriptionally distinct clusters. Both 15-GEP class and *PRAME* status were associated with shifts in the relative abundance of transcriptionally defined T cell states.

**Figure 6.**
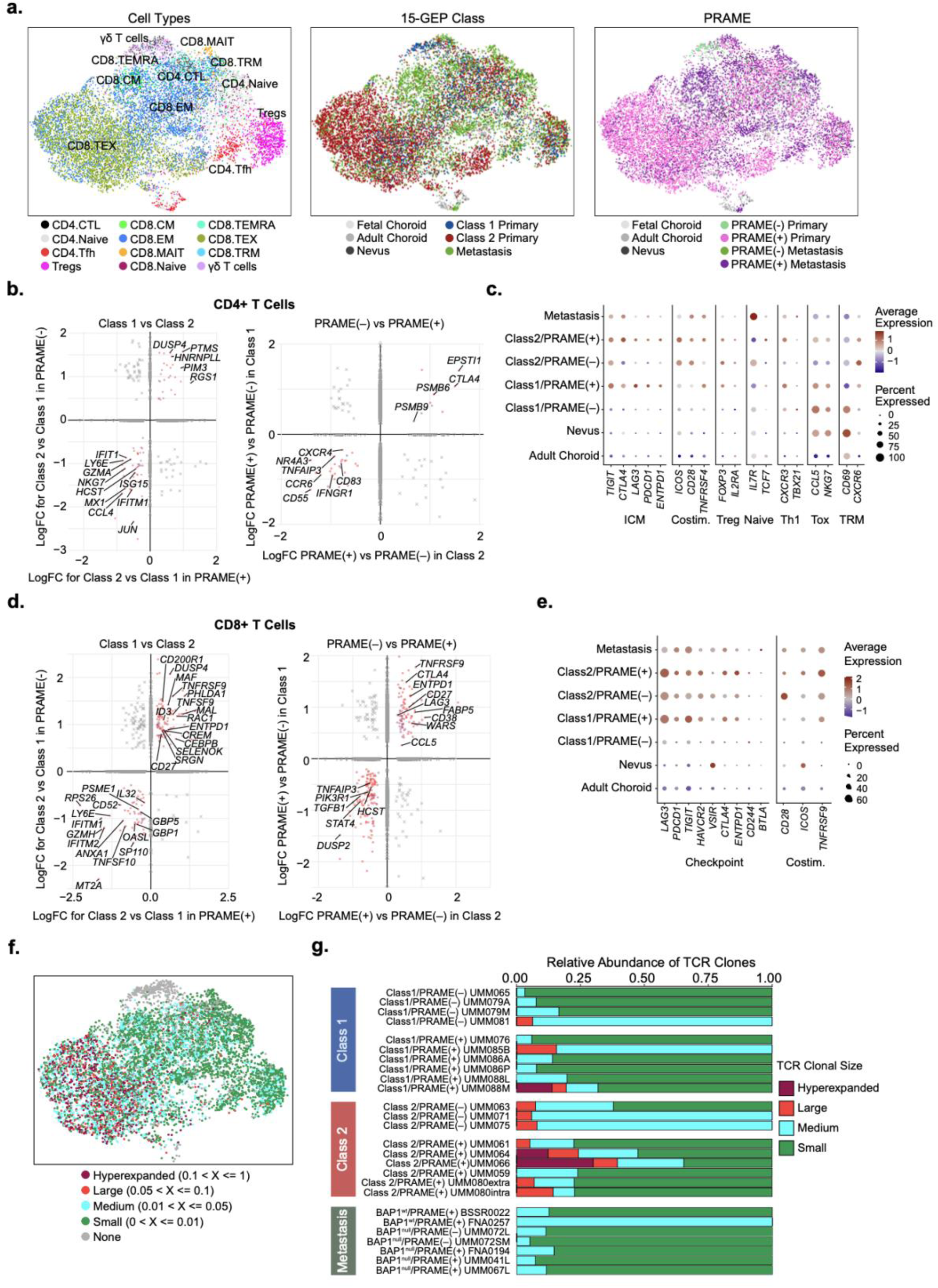
T cell transcriptional programs and subtype composition across uveal melanoma molecular subgroups. **a**, Dimensionality reduction by Uniform Manifold Approximation and Projection (UMAP) plots depicting T cell populations (*n =* 12,266 cells) annotated according to subtype, 15-GEP class and *PRAME* expression status. **b**, Double-volcano plot showing log_2_ fold-change (log_2_FC) of differentially expressed genes (DEGs) in CD4+ T cells from Class 2 versus Class 1 (left) and PRAME(+) versus PRAME(–) (right) primary tumors (FDR-adjusted *P* < 0.05, absolute log2FC > 0.2). In the left volcano plot, the x-axis shows log2FC for the Class 2 versus Class 1 comparison among PRAME(+) tumors, and the y-axis shows log2FC for the same comparison among PRAME(–) tumors. In the right volcano plot, the x-axis and y-axis show log2FC for the PRAME(+) versus PRAME(–) comparison within Class 1 and Class 2 tumors, respectively. Key genes that are mutually upregulated (upper right quadrant) or downregulated (lower left quadrant) are highlighted. Complete DEG analysis results are available in Source Data. **c**, Dot plot of CD4+ T cell functional state and lineage marker genes grouped by immune checkpoint molecule (ICM), costimulatory (Costim.), regulatory T cell (Treg), naïve, Th1, cytotoxic (Tox), and tissue-resident memory (TRM) gene sets, across tumor subtypes. Dot size indicates the percent of cells expressing each gene, and dot color indicates the average normalized expression. **d**, Double-volcano plot showing log_2_FC of DEGs in CD8+ T cells from Class 2 versus Class 1 (left) and PRAME(+) versus PRAME(–) (right) primary tumors (FDR-adjusted *P* < 0.05, absolute log2FC > 0.2). In the left volcano plot, the x-axis and y-axis indicate the log_2_FC for the DEG analysis in PRAME(+) and PRAME(–) primary tumors, respectively. In the right volcano plot, the x-axis and y-axis indicate the log_2_FC for the DEG analysis in Class 2 and Class 1 primary tumors, respectively. Key genes that are mutually upregulated (upper right quadrant) or downregulated (lower left quadrant) are highlighted. Complete DEG analysis results are available in Source Data. **e**, Dot plot of immune checkpoint molecule and costimulatory genes in CD8+ T cells across tumor subtypes. **f**, UMAP of CD8+ T cells annotated by TCR clonal size category, based on scTCR-seq repertoires across 26 samples from 15 primary tumors and 7 metastatic tumors). Clones were classified as hyper-expanded (>10% of the repertoire), large (5–10%), medium (1– 5%) or small (<1% of all TCR clones). T cells with no TCR clones identified were labeled as “None.” **g**, Stacked bar plot of TCR repertoire clone size composition per sample, shown as fraction of total abundance. Samples are grouped by tumor type (Class 1 primary, Class 2 primary, or metastasis) and, within each group, by *PRAME* status; metastatic samples are further annotated by BAP1 status (wild-type or null).

DEG analysis of CD4^+^ and CD8^+^ T cells revealed distinct transcriptional programs associated with both 15-GEP class and *PRAME* status (**Fig. 6b–e**). CD4^+^ T cells from Class 2 tumors exhibited increased expression of genes involved in negative regulation of MAPK and G protein-coupled receptor signaling (e.g., *DUSP4*, *RGS1*), accompanied by reduced expression of interferon-stimulated genes (e.g., *IFIT1*, *IFITM1*, *ISG15*) and cytotoxic/chemokine effector genes (e.g., *GZMA*, *CCL4*), consistent with attenuated signal transduction, interferon responsiveness, and effector activation (**Fig. 6b**). In contrast, CD4^+^ T cells from PRAME(+) tumors exhibited increased expression of genes involved in antigen processing (e.g., *PSMB6*, *PSMB9*), immune checkpoint regulation (e.g., *CTLA4*), and interferon-associated immune activation (e.g., *EPSTI1*), together with reduced expression of genes involved in immune activation and trafficking (e.g., *CXCR4*, *CD83*), interferon responsiveness (e.g., *IFNGR1*), and complement regulation (e.g., *CD55*), consistent with enhanced immune checkpoint regulation, altered antigen-processing capacity, and diminished immune responsiveness (**Fig. 6b**). Analysis of representative CD4^+^ T cell functional and lineage markers revealed coordinated upregulation of checkpoint molecules in PRAME(+) primary tumors, whereas their expression was minimal in adult choroid and nevi and generally lower in PRAME(–) tumors (**Fig. 6c**). Costimulatory, Treg and Th1 markers varied across subgroups, while cytotoxic and tissue-resident programs were most prominent in nevi and Class 1/PRAME(–) tumors and generally diminished in higher-risk primary and metastatic tumors. Metastatic CD4^+^ T cells were distinguished by prominent IL7R expression.

CD8^+^ T cells from Class 2 tumors exhibited increased expression of genes involved in costimulation (e.g., *TNFRSF9*, *CD27*), immune regulation (e.g., *ENTPD1*, *CD200R1*) and T cell activation (e.g., *DUSP4*, *MAF*), together with reduced expression of genes involved in cytotoxic effector function (e.g., *GZMH*, *TNFSF10*, *IL32*) and interferon responses (e.g., *IFITM1*, *IFITM2*, *OASL*, *GBP1*, *GBP5*)(**Fig. 6d**). Similarly, CD8^+^ T cells from PRAME(+) tumors exhibited increased expression of genes involved in T-cell activation (e.g., *CD38*, *CCL5*), costimulation (e.g., *TNFRSF9*, *CD27*), and immune regulation (e.g., *ENTPD1*), together with reduced expression of genes involved in T cell signaling and homeostasis (e.g., *DUSP2*, *STAT4*, *PIK3R1*, *TNFAIP3*) (**Fig. 6d**).

For CD8^+^ T cells, a curated panel set of representative costimulatory and checkpoint genes demonstrated progressive, coordinated upregulation of costimulatory and checkpoint molecules during malignant evolution (**Fig. 6e** and **Extended Data Fig. 4a**). Compared with normal adult uveal and uveal nevus samples, which exhibited minimal expression of these receptors, Class 1/PRAME(–) tumors showed the emergence of a coordinated activation regulatory program characterized by simultaneous induction of costimulatory (*CD28*, *TNFRSF9*) and checkpoint (*LAG3*, *PDCD1*, *TIGIT*, *HAVCR2*, *CTLA4*, *ENTPD1*) receptors, suggesting that remodeling of the adaptive immune microenvironment accompanies malignant evolution. The CD8^+^ T cell compartment continued to evolve to a chronically activated, checkpoint-rich phenotype accompanying molecular progression from Class 1/PRAME(+) through Class 2/PRAME(+) tumors. Metastatic tumors generally retained this phenotype, although *LAG3* expression tended to be reduced compared to Class 2 primary tumors. These checkpoint and costimulatory expression patterns were highly concordant within each molecular subgroup and were remarkably similar among anatomically distinct regions sampled from the same tumor. Together, these findings indicate that Class 2 and *PRAME* expression exert distinct but complementary effects on CD4^+^ T cell transcriptional programs, whereas they converge on a common activated, checkpoint- and costimulatory receptor-rich CD8^+^ T cell phenotype with attenuated cytotoxic and interferon-responsive programs.

To further investigate the relationship between CD8^+^ T cell states and antigen recognition, we performed single-cell TCR sequencing on 26 samples from 22 tumors (**Fig. 6f-g** and **Extended Data Fig. 4b**). Hyperexpanded and large T cell clones localized predominantly within chronically activated, checkpoint-enriched CD8^+^ T cell populations from Class 2 and PRAME(+) primary tumors, indicating persistent antigen-driven clonal expansion despite concurrent upregulation of immune checkpoint pathways. Notably, the absence of large and hyperexpanded clones in metastatic tumors suggests contraction or remodeling of the T cell repertoire during metastatic progression, perhaps through immune editing or altered antigenic stimulation in the metastatic microenvironment. Multiregional analysis demonstrated extensive sharing of expanded T cell clonotypes across spatially distinct regions of the same tumors, while each region also contained a small subset of uniquely expanded clonotypes (**Extended Data Fig. 4b**), suggesting that the dominant intratumoral T cell repertoire is broadly conserved while continuing to diversify. In one case, the UM had undergone extraocular invasion into the orbit prior to surgical excision by enucleation, and we were able to obtain intraocular and extraocular samples for analysis (UMM080intra and UMM080extra, respectively). Despite being sampled from anatomically distinct sites that likely represent different immune microenvironments, most expanded clonotypes were shared between the two samples. In another case, asynchronous metastases to liver and skin (UMM072L and UMM072SM) were sampled and analyzed. Both metastatic lesions retained a shared core of pre-existing expanded clonotypes, with additional site-specific clonotypes emerging later within each metastatic niche. Collectively, these findings indicate that BAP1 loss and *PRAME* expression are associated with coordinated remodeling of the adaptive T cell compartment, characterized by a shift from cytotoxic, interferon-responsive states toward chronically activated, antigen-experienced populations enriched for checkpoint and costimulatory molecules.

### *PRAME* expression is associated with a shift in NK cell composition with minimal transcriptional reprogramming

Given the profound adaptive immune remodeling associated with BAP1 loss and *PRAME* expression, we next examined whether analogous transcriptional changes occurred within the NK cell compartment. A total of 321 NK cells were identified across all tumor samples, among which 5 main transcriptional subtypes were identified and annotated according to their dominant marker expression, including CD16^hi^CD56^lo^, CD16^lo^CD11B^hi^, CD16^lo^CD27^hi^, CD16^lo^CD56^hi^ and CD16^lo^CD69^hi^ (**Fig. 7a**). Whereas there was no significant clustering of NK cell subtypes according to 15-GEP class, *PRAME* expression was associated with a shift in subset composition characterized by relative enrichment of CD16^hi^CD56^lo^ NK cells and a corresponding reduction in CD16^lo^CD69^hi^ NK cells (**Fig. 7b**). We then examined the expression of representative NK cell activation/effector, inhibitory/regulatory, and lineage-associated genes across individual specimens (**Fig. 7c**). In contrast to the marked transcriptional changes observed in CD4^+^ and CD8^+^ T cells, NK cells demonstrated remarkably stable transcriptional programs across normal uvea, uveal nevus, primary and metastatic tumor samples. Canonical activation and effector genes (e.g., *IFNG*, *IL32*, *XCL1*, *XCL2*, *JUN*, *JUND*, *FOS*, *NCR1*, *NCR3*, *KLRK1*, *TNFRSF18*), remained broadly conserved across molecular subgroups, while inhibitory receptors and immune regulatory molecules (e.g., *KLRC1*, *HAVCR2*, *TIGIT*, *VSIR*, *LAG3*, *BTLA*, *CTLA4*, *PDCD1*, *CISH*, *IDO1*, *PTGER4*, *ADORA2A*) exhibited only modest intertumoral variability without coordinated upregulation associated with 15-GEP class or *PRAME* status or with metastatic status. Likewise, lineage-associated markers (*CD44* and *CXCR4*) remained largely unchanged. Interestingly, NK cells from normal adult choroid exhibited a more activated transcriptional program than NK cells from uveal nevi and melanoma samples, possibly representing a poised surveillance state in this immune-privileged tissue which does not persist with neoplastic progression. Together, these findings indicate that, unlike the profound adaptive immune remodeling observed in macrophages and T cells, the NK cell compartment remained comparatively stable with *PRAME* expression being associated with a shift from tissue-associated activated NK cells to more mature circulating-type cytotoxic NK cells.

**Figure 7.**
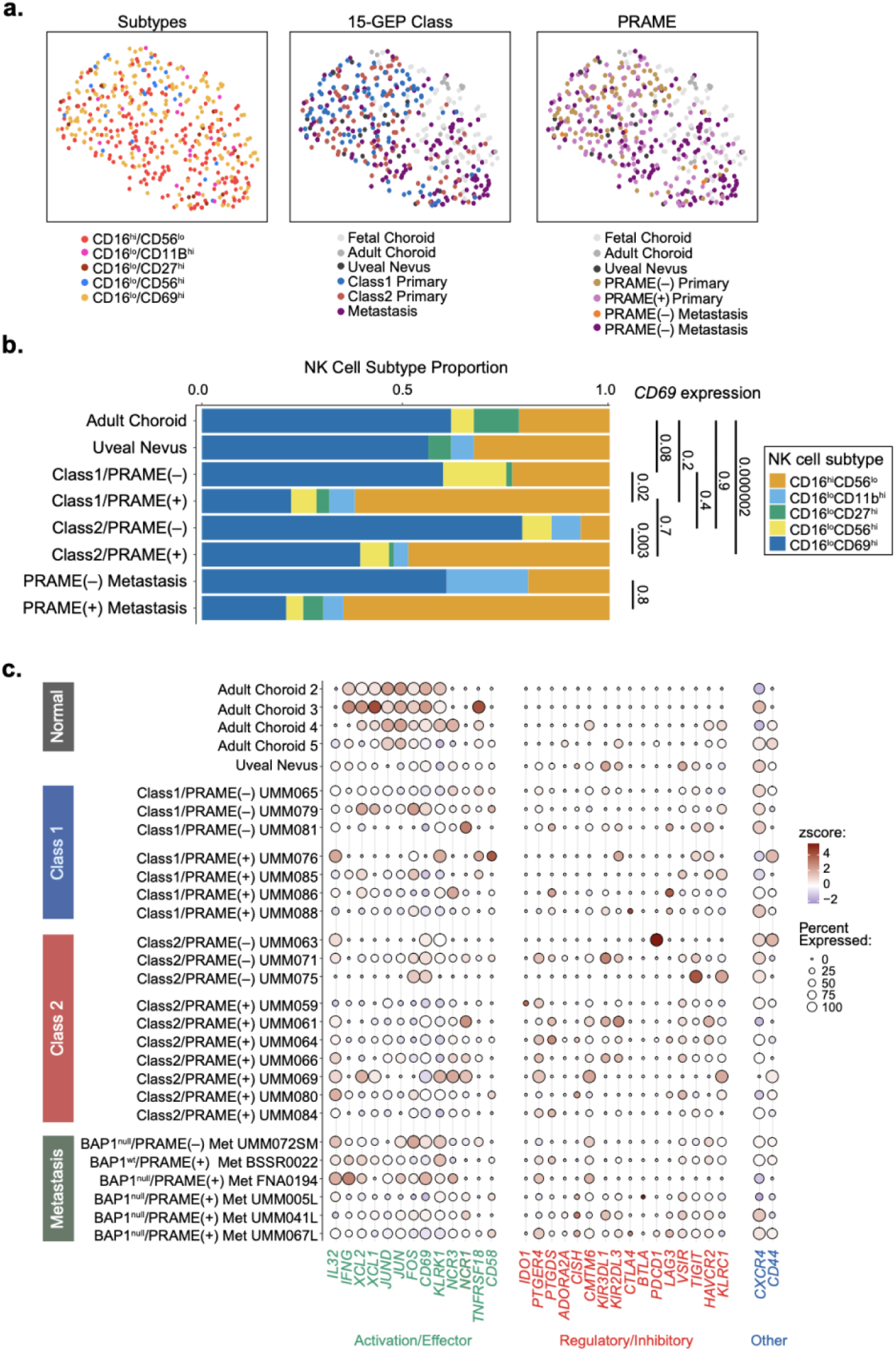
NK cell subtype composition across uveal melanoma molecular subgroups. **a**, Uniform Manifold Approximation and Projection (UMAP) plots of NK cells (*n =* 435) colored according to NK cell subtype (left), 15-GEP class (middle) and *PRAME* status (right). **b**, Stacked bar plot showing the relative proportions of five NK cell subtypes in samples from adult choroid (*n* = 18), uveal nevus (*n* = 18), primary UM (grouped by 15-GEP class and *PRAME* expression status) (*n* = 196) and metastatic UM (grouped by *PRAME* expression status) (*n* = 125). NK cells were classified into marker-defined states, labeled CD16^hi^CD56^lo^ by default unless normalized expression of CD56, CD69, CD27 or CD11b (ITGAM) exceeded that of CD16, in which case cells were classified as CD16^lo^CD56^hi^, CD16^lo^CD69^hi^, CD16^lo^CD27^hi^ or CD16^lo^CD11b^hi^, respectively. FDR-adjusted *P* values for pairwise comparisons of *CD69* expression between the indicated groups are shown on the right. **c,** Dot plot showing the expression of 13 activating/effector genes (green), 16 regulatory/inhibitory genes (red) and 2 additional immune-regulatory genes (blue) across individual samples grouped according to genomic subtype as indicated. Dot size denotes the percentage of NK cells expressing each gene, and color denotes scaled mean expression (z-score).

### PRAME-enriched tumor regions form spatially organized immunoregulatory niches

Having identified *PRAME* expression in UM cells as a key organizing axis of malignant evolution associated with immunoregulatory remodeling across tumor and immune compartments, we next asked whether these transcriptional changes converge within spatially organized intratumoral niches. To examine PRAME-associated spatial organization independently of BAP1 loss, we performed spatial transcriptomic profiling of three Class 1 primary UM samples containing minimal (UMM079), low (UMM085) and high (UMM086) proportions of *PRAME*-expressing tumor cells (**Extended Data Fig. 5a-f**). Cell type annotations were derived from matched scRNA-seq data from the same tumors and were projected onto spatial capture spots, enabling spatial mapping of cell populations.

We first examined UMM086, which contained the highest proportion of PRAME(+) tumor cells (**Fig. 8a-b**). Spatial mapping revealed marked intratumoral heterogeneity in both PRAME(+) tumor cells and cell type-enriched capture spots. PRAME(+) regions were enriched in discrete areas throughout the tumor and were spatially associated with specific immune cell populations. To identify biological programs that may be spatially associated with PRAME(+) regions, we correlated the expression of *PRAME* with that of all genes expressed in each capture spot, ranked genes according to their correlation with *PRAME* expression across all spots, and performed GSEA of the resulting ranked gene list (**Fig. 8c**). *PRAME*-positive regions were significantly enriched for pathways involved in the MHC protein complex, leukocyte-mediated cytotoxicity, DNA damage response, B cell-mediated immunity and interferon signaling. To quantify these spatial relationships, we performed proximity enrichment analysis across inferred cell populations (**Fig. 8d**). B lineage cells and PRAME(+) UM cells showed strong homotypic enrichment, consistent with spatial clustering. Among heterotypic interactions, B lineage cells were enriched near macrophages, and both immune populations were preferentially associated with PRAME(+) UM cells. By contrast, macrophages and B lineage cells were depleted near the broader UM cell population, while PRAME(+) UM cells were depleted in proximity to other UM cells. These findings support the presence of spatially discrete PRAME(+) tumor–immune niches enriched for macrophages and B lineage cells. Building on these proximity-enrichment findings, we further examined the spatial relationships among PRAME(+) UM cells, macrophages and B lineage cells using complementary colocalization analyses (**Fig. 8e–g**). Pairwise spatial overlays confirmed that macrophages and B lineage cells frequently occupied the same discrete regions as PRAME(+) UM cells and colocalized with one another (**Fig. 8e**). Density-based analysis in UMAP space similarly identified concentrated regions of colocalization between PRAME(+) UM cells and both macrophages and B lineage cells, whereas colocalization with T cells was comparatively weaker (**Fig. 8f**). Consistent with these findings, *PRAME* expression was positively correlated with macrophage and B lineage cell signature scores across capture spots, while its correlation with the T cell signature was substantially weaker (**Fig. 8g**). Together, these complementary analyses corroborated the preferential spatial association of PRAME(+) UM cells with macrophages and B lineage cells and further defined these populations as prominent components of localized PRAME-associated tumor–immune niches.

**Figure 8.**
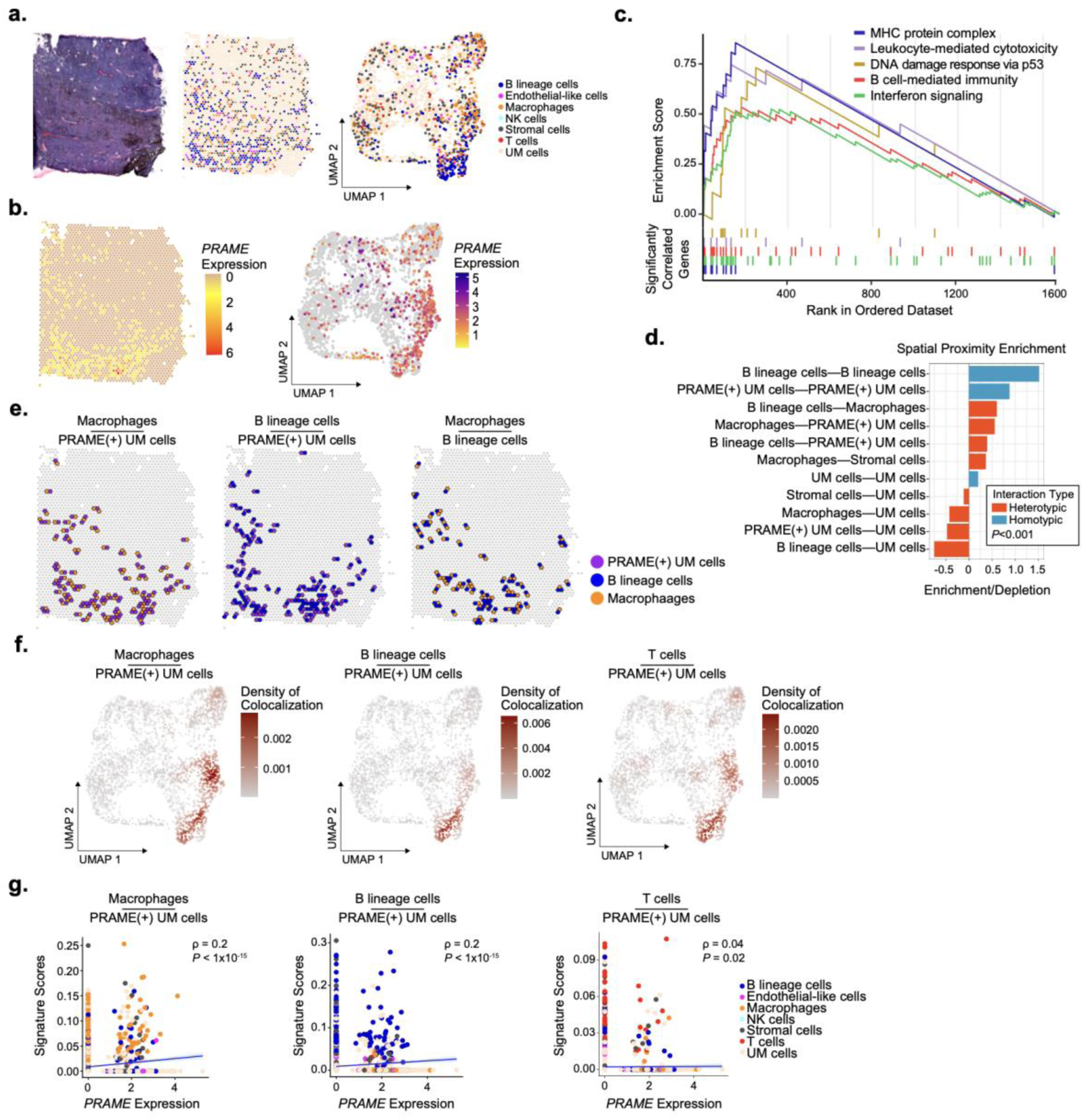
Spatial organization of immune niches in PRAME-enriched uveal melanoma regions. **a**, Tumor section stained with hematoxylin and eosin (H&E; left), 10x Genomics Visium spatial transcriptomic map of capture spots (middle), and uniform manifold approximation and projection (UMAP) embedding of the capture spots (right) from the Class 1/PRAME(+) primary UM sample UMM086. Capture spots (*n* = 2,750) in the middle and right panels are colored by inferred cell type annotation determined by marker-based module scoring with UCell. **b,** Spatial map (left) and UMAP embedding (right) of spatial capture spots colored by normalized *PRAME* expression. **c,** Gene set enrichment analysis (GSEA) of genes ranked by their spatial correlation with *PRAME* expression across Visium capture spots, showing significantly enriched pathways. Detailed summary of complete GSEA results provided in Source Data. **d,** Bar plot showing significantly enriched or depleted cell-type pairs identified by spatial proximity analysis. Positive and negative values indicate spatial enrichment and depletion, respectively. Bar colors distinguish heterotypic (red) and homotypic (blue) pairs. Visium capture spots were annotated using UCell scores for cell type marker modules. Significance was assessed using 10,000 simulations, and all displayed cell-type pairs demonstrated *P* < 0.001. **e,** Spatial maps highlighting colocalization of PRAME(+) UM cells (purple) with macrophages (orange; left) and B lineage cells (blue; middle), and of macrophages with B lineage cells (right). **f,** UMAP embeddings of Visium capture spots colored by the joint density of *PRAME* expression and macrophage (left), B lineage cell (middle) or T cell (right) signature scores. **g,** Scatter plots corresponding to **f**, showing the relationships between normalized *PRAME* expression and macrophage (left), B lineage cell (middle) or T cell (right) signature scores across Visium capture spots. Each point represents one capture spot and is colored by inferred cell type. Spearman rank correlation coefficients (ρ) and corresponding two-sided *P* values are shown. Blue lines indicate linear regression fits and shaded bands denote 95% confidence intervals.

Given the seven recurrent tumor cell transcriptional states identified in our scRNA-seq analyses (**Fig. 2**), we examined how these states may be spatially organized across the three tumors (**Extended Data Fig. 6a-c**). In the PRAME(–) UMM079 sample, the differentiated state predominated throughout the tumor, whereas the proliferative, hypoxic-glycolytic, neural crest-like, stress-responsive and immune-interacting states were sparse or confined to discrete regions (**Extended Data Fig. 6a**). The immune-interacting state was particularly restricted, with its strongest activity localized near the interface between the tumor and adjacent tissue.

UMM085, which contained a small proportion of *PRAME*-expressing tumor cells, similarly retained a broadly distributed differentiated program, with *PRAME* expression and neural crest-like, vasculogenic and immune-interacting activity limited to scattered intratumoral foci (**Extended Data Fig. 6b**). In contrast, UMM086, with a high proportion of *PRAME*-expressing tumor cells, exhibited multiple spatially discrete *PRAME*-enriched regions within a heterogeneous and compartmentalized transcriptional landscape (**Extended Data Fig. 6c**). In contrast to UMM079 and UMM085, the differentiated state in UMM086 was less prominent overall and was particularly diminished within *PRAME*-enriched regions. Neural crest-like, vasculogenic and immune-interacting programs were preferentially associated with PRAME-enriched regions, whereas hypoxic-glycolytic and stress-responsive states were inversely related to *PRAME* expression. The proliferative state remained limited and showed no consistent spatial relationship with *PRAME*. Since our single-cell analyses identified macrophages and B lineage cells as prominent components of the PRAME-associated immune landscape (**Fig. 4b**), we mapped these populations relative to transcriptional states (**Extended Data Fig. 6c**). Spots enriched for macrophage and plasma cell signatures showed substantial spatial concordance and colocalized with regions enriched for vasculogenic and immune-interacting states. Both signatures were inversely associated with the differentiated state and showed weaker inverse relationships with neural crest-like and stress-responsive states, but no clear relationship with proliferative or hypoxic–glycolytic states.

## DISCUSSION

Our findings support a model in which malignant progression is driven by coordinated evolution of malignant cell states and the immune microenvironment along two complementary molecular axes defined by BAP1 loss and *PRAME* expression. Although both alterations are associated with inflammatory and immunoregulatory remodeling, they produce distinct effects across tumor cells, macrophages and CD4^+^ T cells while converging on checkpoint-rich CD8^+^ T cells and HLA-E-associated signaling. These findings extend our understanding of the immunoregulatory consequences of BAP1 loss while repositioning PRAME from a prognostic biomarker and target antigen to a functional contributor to the evolving tumor–immune ecosystem.

A point of convergence between BAP1 loss and *PRAME* expression was the induction of multiple MHC class I genes, most notably HLA-E. In isogenic models, BAP1 depletion and *PRAME* induction each activated interferon and TNF–NF-κB programs and induced multiple MHC class I genes, with HLA-E showing more consistent induction than the classical HLA genes. The inferred HLA-E–CD94/NKG2A interactions suggest that this HLA-E induction may engage an inhibitory checkpoint on NK and CD8^+^ T cells, allowing inflamed MHC class I-high tumors to resist immune elimination. Direct functional studies will be needed to test this hypothesis. More broadly, these findings highlight a paradox that inflammatory and antigen presentation capacity need not translate into effective antitumor immunity but can be co-opted to drive immune suppression and tumor progression.

Spatial profiling revealed that *PRAME*-expressing tumor cells formed nonrandom clusters within regions preferentially associated with neural crest-like, vasculogenic and immune-interacting tumor states and with macrophage and plasma cell signatures. Their association with T cells was substantially weaker. These regions were also enriched for MHC, interferon and B cell immunity programs and showed relative depletion of the differentiated tumor state. Together, these findings link PRAME to a spatially restricted tissue ecology coupling tumor cell dedifferentiation and inflammatory signaling to niches enriched for macrophages and plasma cells. The presence of this organization within a Class 1 background implicates PRAME as a potential organizer of the TIM independently of BAP1 loss.

The putative transcriptional continuum extending from normal melanocytes and nevus cells through the molecular UM subgroups placed progressive loss of differentiated identity at the center of malignant evolution. Rather than generating wholly new phenotypes, advancing tumors increasingly deployed a limited repertoire of neural crest-like, stress-responsive, hypoxic-glycolytic and immune-interacting states at the expense of depletion of tumor cells with the differentiated state. Metastatic cells recapitulated the same states observed in primary tumors, but with altered abundances and no detectable metastasis-specific state. Together, these findings support a model in which plasticity generates recurrent cell states while selection favors their context-dependent expansion and spatial organization. Metastatic competence may therefore arise through altered deployment and organization of pre-existing cell states rather than acquisition of a single metastasis-specific transcriptional program.

T cell evolution in UM was characterized not only by increasing exhaustion but also by progressive attenuation of antitumor function despite persistent tumor recognition. Checkpoint and costimulatory receptors increased in parallel across advancing disease stages, while expanded clonotypes were concentrated within chronically activated CD8^+^ T cell populations. The sharing of clonotypes across spatially separated tumor regions was consistent with recognition of antigens distributed throughout the tumor, whereas site-restricted clonotypes indicated continued local diversification of the repertoire. By contrast, the absence of large or hyperexpanded clonotypes in metastases, despite their presence in high-risk primary tumors, suggested repertoire contraction or remodeling during dissemination. Together, these findings support a model of cancer immune evolution in which antigen recognition persists but becomes progressively constrained by chronic stimulation and immunoregulatory circuitry.

These findings raise several testable therapeutic hypotheses. The convergence of BAP1 loss and *PRAME* expression on HLA-E nominates the HLA-E–CD94/NKG2A axis as a candidate immunoregulatory vulnerability in high-risk UM. PRAME may therefore represent both a therapeutic antigen and a source of immunoregulatory programs that could constrain PRAME-directed immunity, suggesting that the efficacy of PRAME-targeted therapies may depend on counteracting these programs. Moreover, the spatial association of *PRAME*-expressing tumor cells with macrophages and plasma cells raises the possibility that effective therapy will require simultaneous targeting of malignant cells and their local multicellular ecosystem.

Several limitations of this study identify priorities for further investigation. The putative transcriptional continuum was inferred from cross-sectional data from different patients and therefore does not establish temporal progression, while inclusion of only one uveal nevus limits conclusions about premalignant evolution. Spatial profiling was restricted to three Class 1 tumors, with the clearest niche architecture observed in one PRAME-high tumor, requiring validation in larger cohorts. CellChat interactions and HLA-E-mediated immune suppression were inferred and require experimental validation, whereas the small number of NK cells limited analysis of this compartment. Despite these limitations, our findings show how tumor cell evolution can reorganize a conserved repertoire of malignant states while simultaneously reshaping the composition, functional state and spatial architecture of the immune microenvironment. More broadly, they reveal how tumor cell plasticity is coordinated with localized immune regulation during cancer evolution and nominate PRAME and HLA-E as potential points of therapeutic intervention.

## EXTENDED DATA FIGURES

**Extended Data Fig. 1.**
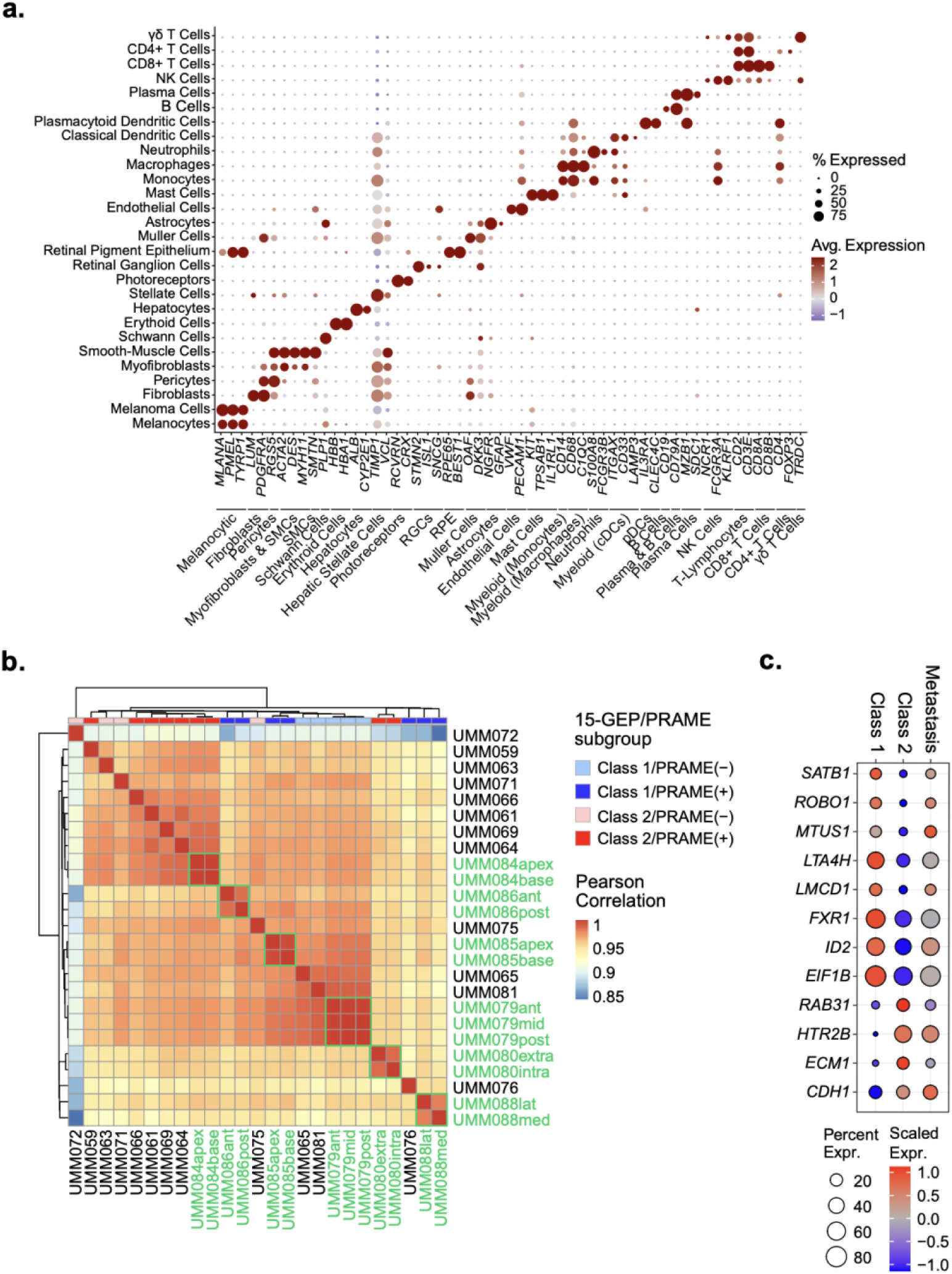
Cell type annotation and transcriptional features of the uveal melanoma single-cell cohort. **a,** Dot plot showing the expression of canonical marker genes used to annotate major cell populations in the integrated single-cell RNA-sequencing dataset of 190,535 cells. Dot size denotes the percentage of cells expressing each gene, and color denotes scaled mean expression within each cell type. **b,** Heatmap of pairwise Pearson correlation coefficients between sample-level pseudobulk transcriptomic profiles generated by sum aggregation of log-normalized RNA values. Color annotations indicate molecular subgroups based on 15-gene expression profile (15-GEP) class and *PRAME* expression status. Green labels identify multiregional samples from a single tumor, and green boxes indicate within-tumor comparisons. **c,** Dot plot showing expression of the 12 differentially expressed prognostic genes from the 15-GEP in UM cells from Class 1 primary, Class 2 primary and metastatic tumors. Dot size denotes the percentage of cells expressing each gene and color denotes scaled mean expression within each group. Raw data are available in Source Data.

**Extended Data Fig. 2.**
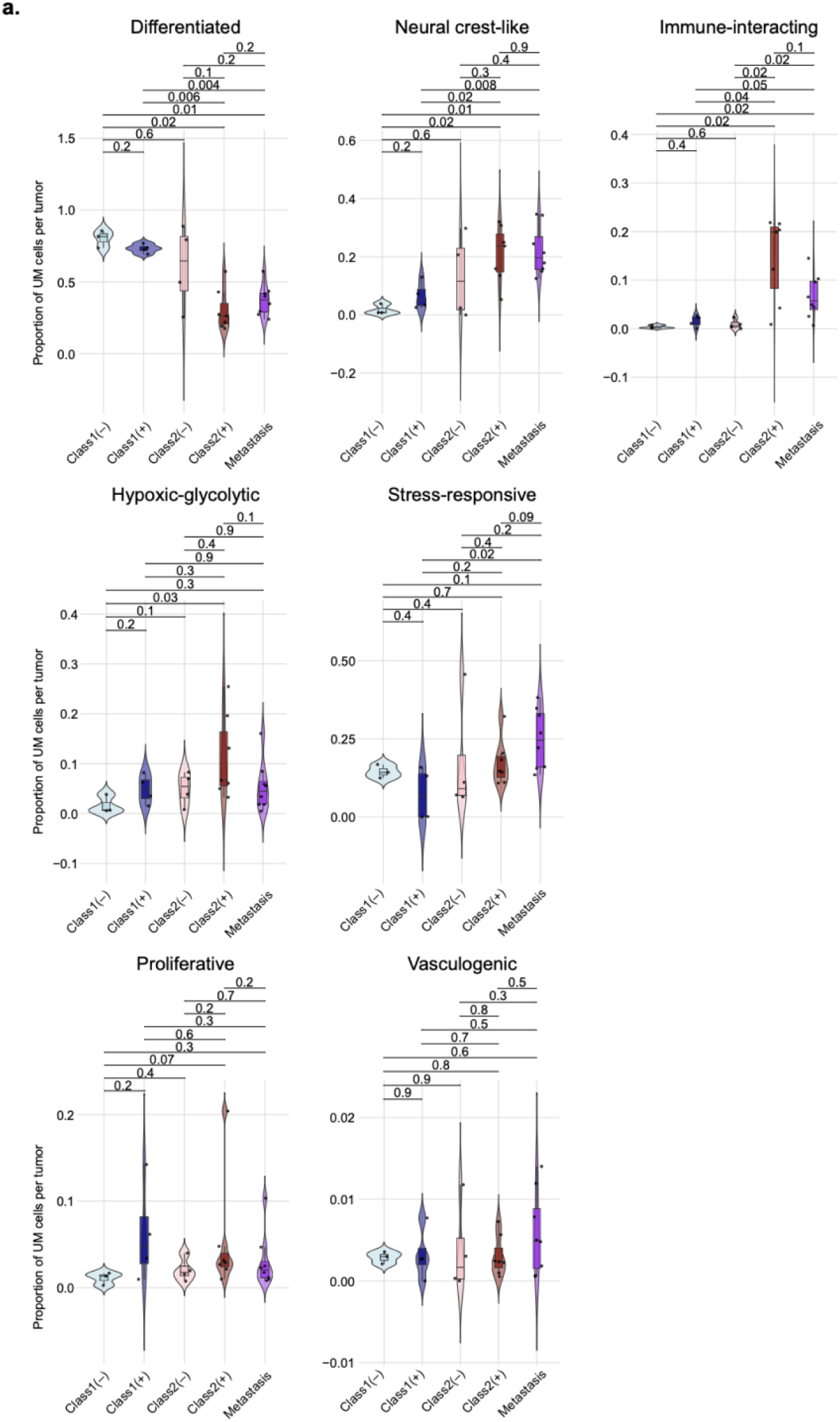
Distribution of uveal melanoma transcriptional states across molecular subgroups. Violin plots with embedded box-and-whisker plots demonstrating the abundance of uveal melanoma (UM) cells assigned to each transcriptional state for each tumor sample (represented by dots). Tumor abundances were grouped according to genomic subtype, including Class 1/PRAME(−) (*n* = 3), Class 1/PRAME(+) (*n* = 4), Class 2/PRAME(−) (*n* = 4) and Class 2/PRAME(+) (*n* = 7) primary tumors and metastases (*n* = 8). Boxes show the median and interquartile range (IQR), and whiskers extend to the most extreme values within 1.5 × IQR. Violins were scaled to ensure all had equal total area. Pairwise significance testing was performed using two-sided Wilcoxon rank-sum tests. Raw data used to generate plots are available in Source Data.

**Extended Data Fig. 3.**
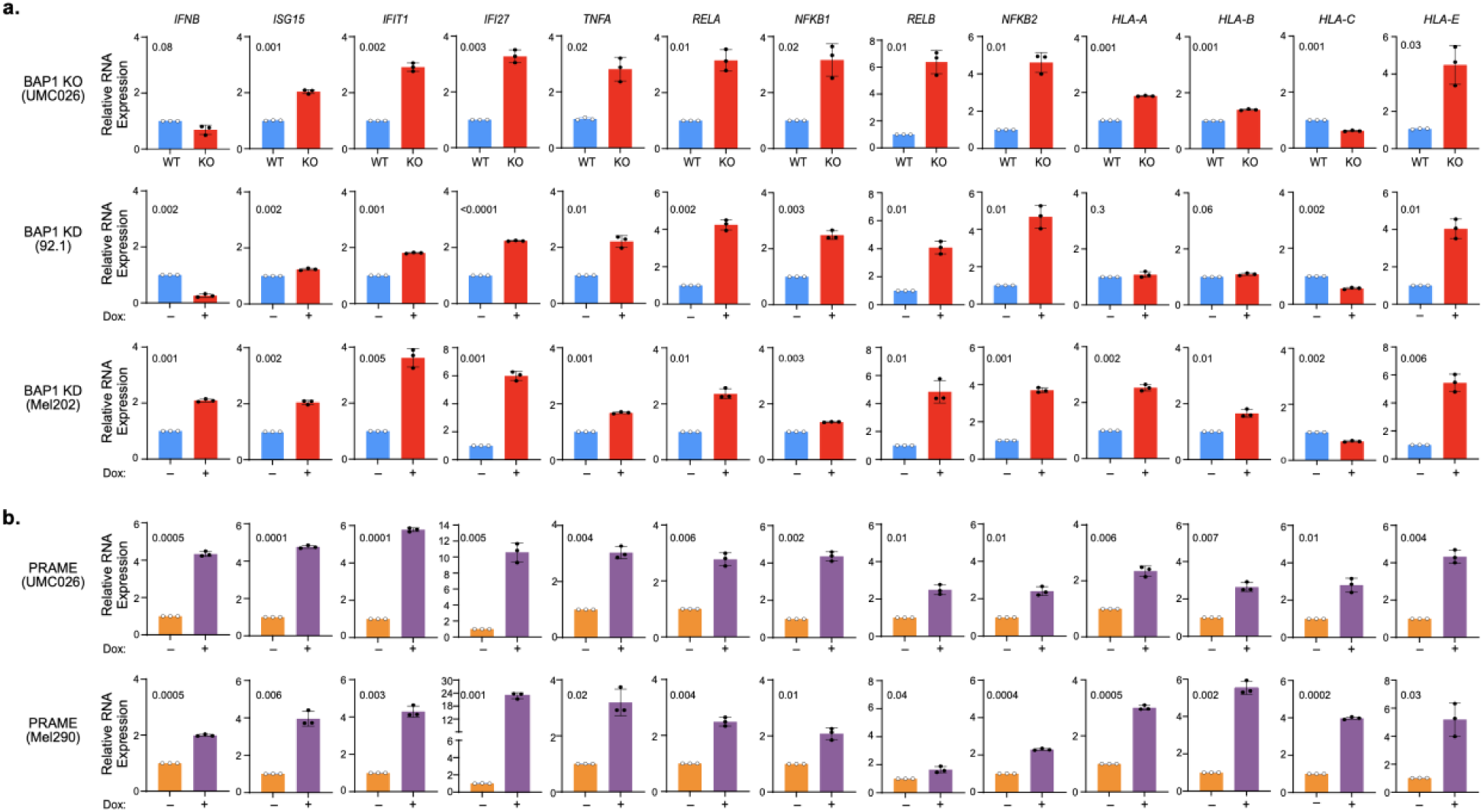
BAP1 loss and *PRAME* expression alter type I interferon, NF-κB and MHC class I gene expression in cells of uveal melanocytic lineage. **a,** Reverse-transcription quantitative PCR (RT–qPCR) analysis of type I interferon-responsive (*IFNB1, ISG15, IFIT1* and *IFI27*), NF-κB-associated (*TNF, RELA, NFKB1, RELB* and *NFKB2*) and MHC class I (*HLA-A, HLA-B, HLA-C* and *HLA-E*) transcripts in genetically engineered uveal melanocytic models of BAP1 loss. CRISPR–Cas9-generated *BAP1* knockout (KO) UMC026 nonmalignant uveal melanocytes were compared with wild-type (WT) controls. The 92.1 and Mel202 UM cell lines containing a tetracycline-inducible shRNA targeting *BAP1* were cultured without (Dox−) or with (Dox+) doxycycline to induce *BAP1* knockdown (KD). Blue bars denote BAP1-intact controls and red bars denote BAP1-deficient conditions. **b,** RT–qPCR analysis of the same transcripts in UMC026 and Mel290 cells engineered with tetracycline-inducible *PRAME* expression and cultured without (Dox−; orange) or with (Dox+; purple) doxycycline. For **a,b**, transcript abundance was normalized to 18S rRNA and plotted as fold change relative to the corresponding control condition. Bars show the mean of *n* = 3 independent biological replicates, points denote individual replicates, and error bars show standard deviation. *P* values were calculated using two-sided unpaired Student’s *t*-tests with Welch’s correction and are shown above each comparison. Raw values are available in Source Data.

**Extended Data Fig. 4.**
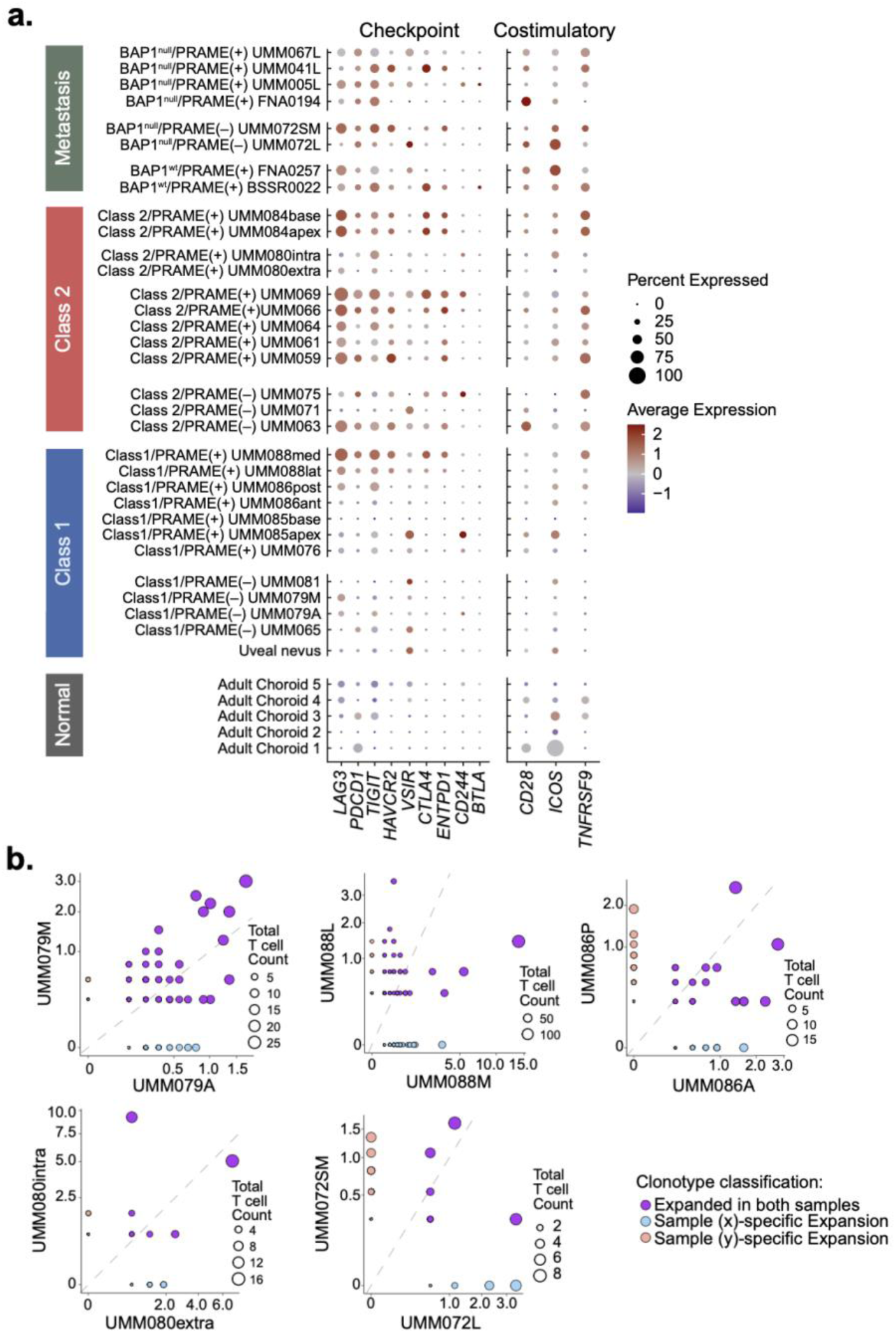
CD8^+^ T cell checkpoint and costimulatory gene expression and TCR clonotype sharing in uveal melanoma. **a**, Dot plot showing checkpoint and costimulatory gene expression in CD8^+^ T cells across individual single-cell RNA sequencing (scRNA-seq) profiles. Samples are grouped by tissue or disease category, 15-gene expression profile (15-GEP) class and *PRAME* status. Primary tumors are annotated by 15-GEP class (Class 1 or Class 2) and *PRAME* expression status (negative or positive), whereas metastases are annotated by BAP1 mutation status (BAP1^null^ or BAP1^WT^) and *PRAME* expression status (negative or positive). Dot size denotes the percentage of CD8^+^ T cells expressing each gene and color denotes scaled mean expression. **b**, Bubble plots comparing T cell receptor (TCR) clonotype distributions between paired regions of multiregional primary tumors or paired metastatic sites from the same patient, as determined by single-cell TCR sequencing (scTCR-seq). Each bubble represents a TCR clonotype; bubble size denotes the total number of T cells assigned to that clonotype across the paired samples, and the x and y axes show its relative abundance within the indicated samples. The dashed diagonal denotes equal relative abundance in the paired samples. Blue and orange bubbles indicate clonotypes expanded specifically in the x-axis and y-axis samples, respectively, whereas purple bubbles indicate clonotypes expanded in both samples.

**Extended Data Fig. 5.**
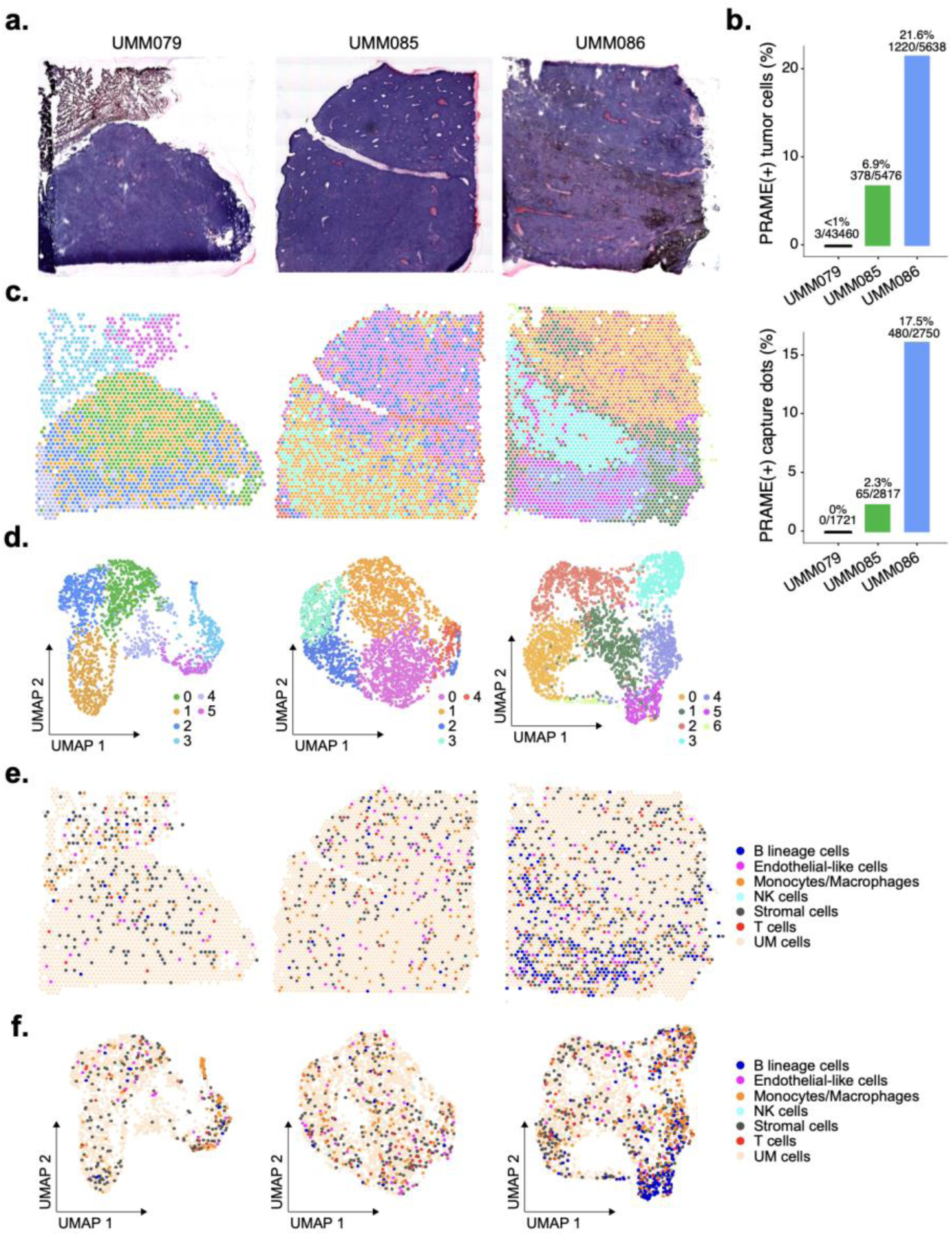
Spatial transcriptomic profiling of Class 1 uveal melanomas with differing *PRAME* expression. **a**, H&E-stained sections from three primary Class 1 uveal melanomas (UMs) analyzed by 10x Genomics Visium spatial transcriptomics: UMM079 (left), UMM085 (middle) and UMM086 (right). **b**, Percentage of UM cells expressing *PRAME* determined by single-cell RNA sequencing (scRNA-seq) (top) and percentage of spatial capture spots expressing *PRAME* (bottom) for UMM079, UMM085 and UMM086. The scRNA-seq analysis included 43,460, 5,476 and 5,638 UM cells, respectively, and the spatial transcriptomic analysis included 1,721, 2,817 and 2,750 capture spots, respectively. **c**, Spatial maps of capture spots from UMM079, UMM085 and UMM086 (left to right) colored by Louvain cluster assignment. **d**, Uniform Manifold Approximation and Projection (UMAP) plots of spatial transcriptional profiles from UMM079, UMM085 and UMM086 (left to right) colored by Louvain cluster assignment corresponding to the colors in panel **c**. Each point represents one spatial capture spot. Clustering was performed using the first 7, 5 and 10 principal components, respectively, as determined to be optimal by elbow plot assessment. **e**, Spatial maps of UMM079, UMM085 and UMM086 (left to right) showing cell type annotations inferred from UCell scores derived from cell type-specific marker gene sets. Annotations comprise B lineage cells (B cells and plasma cells), endothelial-like cells, monocytes/macrophages, NK cells, stromal cells, T cells and UM cells. Stromal cells comprise fibroblasts, smooth muscle cells, myofibroblasts and pericytes, whereas T cells comprise CD4^+^ T cells, CD8^+^ T cells and regulatory T cells. **f**, UMAP plots of spatial capture spots from UMM079, UMM085 and UMM086 (left to right) colored by inferred cell type annotation, with colors corresponding to those in panel **e**. See Methods for additional details.

**Extended Data Fig. 6.**
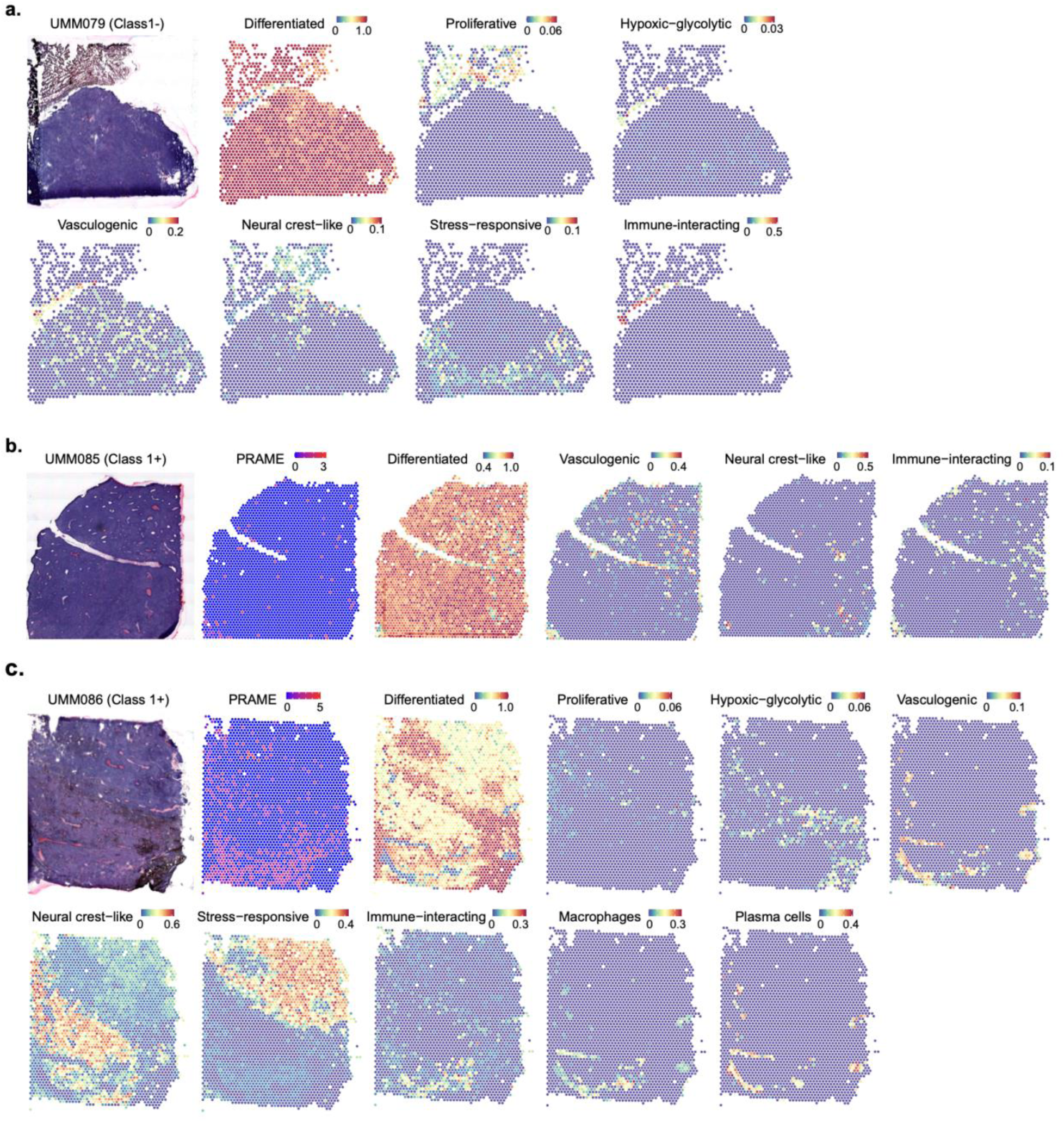
Spatial distribution of tumor cell transcriptional states across Class 1 uveal melanomas with differing *PRAME* expression. **a**, H&E-stained section (left) and spatial maps of UCell signature scores for differentiated, proliferative, hypoxic–glycolytic, vasculogenic, neural crest-like, stress-responsive and immune-interacting tumor cell states in the Class 1/PRAME(−) primary tumor UMM079 with 1,721 spatial capture dots. No *PRAME* expression map displayed for UMM079 due to lack of expression. **b**, H&E-stained section (left) and spatial maps of normalized *PRAME* RNA expression and UCell signature scores for differentiated, vasculogenic, neural crest-like and immune-interacting tumor cell states in the Class 1/PRAME(+) primary tumor UMM085 with 2,817 spatial capture dots. **c**, H&E-stained section (left) and spatial maps of normalized *PRAME* RNA expression, UCell signature scores for all seven tumor cell states and macrophage and plasma cell signature scores in the Class 1/PRAME(+) primary tumor UMM086 with 2,750 spatial capture dots. In **a–c**, each point represents one Visium capture spot, and color indicates normalized *PRAME* RNA expression or the UCell signature score for the indicated tumor cell state or immune cell population. Raw data available in Source Data.

## METHODS

### Tumor sample collection and processing

Human tissue samples were obtained with patient informed consent and approval of the Institutional Review Board of the University of Miami. Immediately following surgical resection, a sample of the tumor was dissected and prepared for single-cell dissociation. Cases that underwent multi-regional sampling after enucleation were labeled according to the tumor region sampled (e.g., anterior, posterior, base, apex) by the surgeon (J.W.H). The comprehensive single-cell atlas comprised 18 primary and 8 metastatic tumors, including 7 primary and 3 metastatic tumors that were previously published^5^, as well as 1 benign congenital nevus from a patient with ocular melanocytosis (*n =* 1; 1,702 cells). Data from these samples were integrated with previously published scRNA-seq data from 5 normal adult choroid (*n =* 12,394 cells) and 4 fetal choroid (*n =* 33,818 cells) tissue samples ^2^, which were provided by the Joint MRC/Wellcome Trust (MR/R006237/1) Human Developmental Biology Resource under ethics permission 08/H0906/21 and by the Northeast-Newcastle and North Tyneside 1 Research Ethics Committee. Tumor samples underwent standard of care molecular testing with the DecisionDx-UM^®^ and DecisionDx^®^-PRAME tests (Castle Biosciences, Inc.) for 15-GEP class and *PRAME* status, respectively, as previously described^11^.

### Tissue processing for single-cell suspension

Collected samples were immediately transferred to gentleMACS C tubes (Miltenyi Biotec) and suspended in 5 mL of DMEM or RPMI1640 Media with 10% FBS and 400 U/mL of collagenase IV for digestion. These UM samples were processed into single-cell suspension using the protocol described in^5^ with consistency to minimize dissociation-associated artifacts in downstream analysis. Miltenyi Red Blood Cell Lysis was conducted when RBC contamination was notable. Miltenyi Cell Debris removal was performed according to manufacturer recommendations. High cell viability (>75%) was confirmed with ThermoFisher LIVE/DEAD Cell Imaging Assay.

### Single-cell RNA and T cell receptor sequencing

Cell suspensions were analyzed using a Nexcelom K2 digital cell counter to determine viability (AO-PI fluorescence) and total particles (bright field). Samples were captured using a 10X Genomics Chromium single cell controller with Immune Profiling V1.1. Chromium Next GEM Single Cell 5ʹ Library and Gel Bead Kit (v1.1, 16 reactions, PN-1000165) and Chromium Single Cell V(D)J Enrichment kit (for Human T Cell, 96 reactions, PN-1000005) were used to generate gene expression and TCR libraries according to the manufacturer’s protocol. cDNA amplification, gene expression and T cell receptor (TCR) library preparation were carried out according to the manufacturer’s instructions. Libraries were sequenced on a NovaSeq 6000 following 10X Genomics sequencing parameters.

### Data processing, quality assessment and clustering

FASTQ files were generated from raw BCL files with the mkfastq command. Raw gene-barcode matrices aligned with the 10X Genomics GRCh38 human reference (version “2020-A”) were generated with the CellRanger (version 5.0.0) count command. Cellbender (version 0.3.0) was used to remove ambient RNA and call cells from raw count matrices. Filtered matrices were read into Seurat (version 5.0.2) using the Read_CellBender_h5_Mat function from scCustomize (version 2.0.1). Seurat was utilized to log-normalize, scale, and cluster samples before doublet identification. DoubletFinder (version 2.0.4) analysis was conducted on each sample, selecting sample-specific pK’s with a maximum mean-variance normalized bimodality coefficient, assuming a doublet rate of 0.8% per 1000 cells sampled (i.e., a doublet rate of 8.0% for a sample with 10,000 cells). With Seurat, all samples were filtered for cells that expressed more than 500 unique features (genes), had more than 1000 unique molecular identifiers (UMIs), and possessed mitochondrial content less than 10 percent. Samples were integrated using stochastic optimization and deep neural network methodology via scVI integration framework^30^ (scvi-tools, version 1.0.4) with a maximum of 400 epochs for training, running Uniform Manifold Approximation and Projection (UMAP) dimensionality reduction and Louvain clustering (resolution = 1) with the first 10 principal components and using 4000 variable features.

Reintegration of immune subset populations was also accomplished via scVI integration with a maximum of 400 training epochs. Batch-adjustment of Principal Component Analysis (PCA) embeddings was conducted with Harmony (version 1.2.3) for other downstream analyses requiring PCA embeddings. Violin plots, dot plots, and other scRNA-seq visualizations were generated with Seurat, scCustomize, and SeuratExtend (version 1.1.2). Heatmaps were generated with ComplexHeatmap (version 2.18.0).

### Cell type annotations

Using UCell (version 2.10.1)^31^, we calculated cell-wise gene set signature enrichment scores. Enrichment scores represented a normalized U statistics (ranging from 0-1) calculated from the RNA expression of provided signature genes using both positive and negative cell markers (**Supplementary Table 2**). A negative marker weight of 0.75 was specified. Analysis was conducted using a maximum gene rank 4000 before considering a gene essentially not expressed. Cells were labelled according to the highest-scoring cell type signature score, as derived from UCell. Cells with a positive UCell-derived melanocytic signature score were labelled as UM cells and melanocytes. Melanocytic cells were reclustered by scVI integration and Louvain clustering (resolution = 1), and clusters enriched for multiple cell-type markers specific for immune cells (including CD3 and CD8A/B) or stromal cells were excluded from cell state analysis for melanocytes and UM cells.

To refine cell type calls from UCell and to enhance cell subtype identification, reference-based annotation was conducted using a pan-cancer immune atlas^29^, with the FindTransferAnchors function from Seurat using 4000 variable features and the first 40 principal components of log-normalized RNA expression for all immune cells. UCell cell type estimates were also utilized to verify reference-based mapping results and to identify rare cell populations, such as stellate cells and hepatocytes. Final cell type annotation was affirmed by marker expression.

T cells underwent CD4^+^ and CD8^+^ T cell subpopulation identification via reference-based mapping utilizing CD4^+^ and CD8^+^ T cell references from a pan-cancer atlas of tumor-infiltrating T cells^32^. Similarly, T cell subpopulations were identified with FindTransferAnchors, using 400 variable features and the first 30 principal components of the log-normalized RNA expression.

Reference-based mapping subtype annotation was used from either CD4^+^ or CD8^+^ T cell reference based on *CD4*, *CD8A* and *CD8B* expression, where the CD4^+^ T cell-based reference was used if *CD4* RNA expression of a T cell was greater than or equal to the expression of *CD8A* or *CD8B*. T cells identified as exhausted cytotoxic-like CD4^+^ T cells were labelled as regulatory T cells when their UCell regulatory T cell signature was greater than 0.5; otherwise, these cells were considered exhausted CD8^+^ T cells. Cells identified as EMOES+ and GNLY+ cytotoxic-like CD4^+^ T cells were all annotated as cytotoxic-like CD4^+^ T cells for downstream analysis.

### UM transcriptional states analysis

The Seurat object (version 5.4.0) with the originally identified 124,744 UM cells from primary (*n* = 25) and metastatic (*n* = 8) tumors was more stringently filtered to 122,715 UM cells by excluding cells expressing canonical immune-lineage markers or having nonzero immune-cell signature scores to minimize immune-cell contamination. RNA expression values were log-normalized with a scale factor of 10,000. The 3,000 most variable genes were selected using the variance-stabilizing transformation method, scaled, and used for principal-component analysis with 50 components. Sample-associated variation was corrected with Harmony (version 1.2.4) using the original sample identity as the grouping variable. UMAP visualization and shared nearest-neighbor graph construction were performed using the first 30 Harmony dimensions. Graph-based clustering was evaluated across resolutions from 0.2 to 0.8. Cluster stability, sample and disease-stage composition, and marker-gene expression were assessed using clustree (version 0.5.1) diagrams, composition summaries, and Wilcoxon rank-sum marker testing. Clustering was conducted with a resolution of 0.6 and used for annotation of UM transcriptional states. DEGs were identified for each cluster with the Seurat function, FindAllMarkers, using a Wilcoxon rank-sum test (with the settings, “only.pos = TRUE, min.pct = 0.10, logfc.threshold = 0.10”). Candidate markers were further filtered using the following criteria: Bonferroni-adjusted P < 0.05, average log2FC ≥ 0.50, expression in ≥ 25% of cells within the cluster and pct.1 - pct.2 ≥ 0.15. Retained genes were ranked using a composite score based on standardized log2FC (45%), detection specificity (25%), within-cluster prevalence (20%), and adjusted statistical significance (10%). The 20 highest-ranking genes per cluster were reviewed, and states were assigned when a majority of markers supported a coherent biological program. Clusters without a coherent alternative program, aside from their baseline melanocytic signature, were classified as differentiated.

Markers for characterized UM transcriptional states were curated from the DEGs from corresponding cluster analysis, prioritizing significant DEGs with greater state-specificity (expression in ≥ 25% of cells within the cluster and pct.1 - pct.2 ≥ 0.15), and DEGs that ranked among the top 20 markers for state-specific clusters. Of 94 markers for transcriptional states, 89 markers met all significance criteria, and 74 markers ranked among the top 20 DEGs for their cluster. Markers manually curated outside these guidelines included *LTA4H*, *HSPA5*, *PDK1*, *NDUFA4L2*, and *SPAG4*. **Supplementary Table 3** provides the complete marker statistics, ranks, and selection status.

These representative marker gene sets were used to score UM transcriptional state signatures in stringently filtered UM cells using log-normalized expression and the UCell package function, AddModuleScore_UCell, set to a maximum ranking of 1,500 genes.

Transcriptional states were visualized on the Harmony-derived UMAP. Violin plots depict transcriptional state abundances summarized by tumor, with violins scaled to have equal area between groups.

### InferCNV analysis of uveal melanoma cells

Copy-number-associated expression patterns were inferred from raw single-cell RNA-sequencing counts using the R package inferCNV (version v1.24.0) (Tickle et al., 2019). Class 1 and Class 2 UM samples were grouped according to seven assigned transcriptional states: baseline differentiated, hypoxic-glycolytic, immune activated, neural crest-like, proliferative, stress responsive, and vasculogenic. A maximum of 2,500 cells was retained for each transcriptional state. Adult human choroidal melanocytes were used as the reference population. Raw counts were extracted from the Seurat RNA assay, and genes were ordered by genomic position using a filtered hg38 gene-order file. Genes below an average-expression cutoff of 0.1 were removed. inferCNV was run using default settings unless otherwise specified. The expression cutoff was set to 0.1; cells were clustered by their assigned groups, subcluster analysis mode and denoising were enabled. The independently generated Class 1 and Class 2 heatmaps were subsequently assembled into the composite figure.

### Differential abundance and cell composition analysis

Differential abundance of cell types was determined by analyzing cell populations as “neighborhoods” determined from a k-nearest neighbors graph representing the cellular connectivity of primary UM tumors using the R package miloR (version 2.2.0)^19^. The k-nearest neighbor graph was calculated with the buildGraph function from miloR, with 30 (k) nearest-neighbors considered during graph construction and using the Harmony-computed reduced dimensional space with 30 dimensions. Neighborhoods were determined with the makeNhoods function. A fourth of all graph vertices were randomly sampled during neighborhood identification and refinement. Calls were refined by computing median profiles for neighborhoods in the Harmony-dimensional reduction and selecting the nearest vertex.

Neighborhood cell counts per tumor, neighborhood distance, and abstracted neighborhood graphs were generated with the functions countCells, calcNhoodDistance, buildNhoodGraph, respectively. Differential abundance testing for 15-GEP Class and PRAME expression status was conducted on neighborhood counts using Quasi-Likelihood F-test (QLF) implemented by edgeR, with FDR correction performed separately to avoid inappropriate testing comparing neighborhoods with overlapping cells. Neighborhood identities were determined by the annotateNhoods function to label cell type identity, neighborhood cluster, 15-GEP Class and PRAME expression status. Differential abundance results were visualized using the functions plotUMAP, plotDAbeeswarm, and plotNhoodGraph.

For relative abundance analysis, percentage of each cell types comprising each primary tumor were compared by two-tailed Wilcoxon test for significance testing.

### Single-cell differential gene expression analysis

DEGs were identified using scRNA-seq counts and applying edgeR with QLF-based methodology, setting cellular detection rate as a covariate^33,34^. For UM cells, the dataset was downsampled to 1000 cells per sample. Significant differentially expressed genes (DEGs) were defined by false discovery rate (FDR)-corrected *P*-value less than 0.05 and absolute value of log_2_ fold change (log_2_FC) greater than 0.2. Concordant Class 2 DEGs were those consistently upregulated or downregulated in Class 2 tumors relative Class 1 tumors regardless of *PRAME* status, and likewise for Class 1 DEGs. Concordant PRAME(+) DEGs were those consistently upregulated or downregulated in PRAME(+) tumors relative to PRAME(–) tumors regardless of 15-GEP class status, and likewise for PRAME(–) tumors. Double volcano plots were generated from log_2_FC of significant DEGs with the ggplot2 R package (version 4.0.3).

### Single-cell pseudo-bulk analysis

Prior to generating pseudobulk counts from scRNA-seq RNA counts, scatter (version 1.26.1) and scuttle (version 1.8.4) were used to filter low-quality cells (with perCellQCMetrics and isOutlier functions) and genes expressed by fewer than 10 cells. RNA counts were log-normalized (with the scuttle function, logNormCounts), and pseudobulk counts were generated by summing RNA counts for each cell type within each sample. PCA analysis of pseudobulk samples was conducted with DESeq2 (version 1.38.3). Sample-level transcriptomes were compared with Pearson correlation using the cor function from the R package, mosaic (version 1.10.2), and plotted with the pheatmap R package (version 1.0.13) using “ward.D2” methodology for hierarchical clustering.

### Pathway enrichment analysis

All collections from Molecular Signatures Databases (MSigDB, version 2024.1.Hs) pathway gene sets were used as reference gene set for GSEA. Gene set enrichment analysis (GSEA) was conducted with clusterProfiler (version 4.6.2) and fsGSEA (version 1.24.0), providing significant DEGs ranked in descending order according to log_2_ fold-change. Tree plots, dot plots, gene network plots, and other GSEA plots were generated with the R packages clusterProfiler, DOSE (version 3.24.2), and enrichplot (version 1.18.3).

### Single-cell inference of 15-GEP classification

For all UM cells, the 12 prognostic GEP genes were normalized with the UCell function, AddModuleScore, providing RNA data and a maximum rank of 40,000 for increased resolution. The support vector machine (SVM) classifier was trained with 30% of all primary UM cells, randomly selected for the training set. The SVM classifier was trained utilizing the svm function from the R package, e1071 (version 1.7-14), employing a linear kernel. Model fitting was conducted with the R package, rminer (version 1.4.6) with the fit function. Ranked importance of GEP genes in the scGEP classifier was calculated with the Importance function from rminer.

The scGEP classifer was applied to UM cells with the predict function from the R package, stats (version 4.3.2).

### Weighted gene co-expression network analysis

Co-expression network analysis was performed employing the hdWGCNA package (version 0.4.05) implemented within the Seurat framework in R. Analysis was conducted on the processed scRNA-seq dataset including UM cells, adult uveal melanocytes, and fetal uveal melanocytes. Genes for weighted gene co-expression network analysis were selected using the SetupForWGCNA function with the “fraction” method, requiring expression in at least 50% of cells to be retained for downstream analysis. To mitigate sparsity associated with single-cell transcriptomic data, metacells were constructed using the MetacellsByGroups function with Harmony-corrected low-dimensional embeddings, nearest-neighbors parameter set to 30, and the maximum shared cells between metacells set to 10.

Normalized metacell expression matrices were subsequently used for network construction, prepared with the SetDatExpr function using normalized RNA assay data. Soft-thresholding powers were evaluated using the TestSoftPowers function with a signed network topology, and a soft-thresholding power of 10 was selected based on scale-free topology criteria. Weighted gene co-expression networks were constructed using the ConstructNetwork function. Following network generation, expression values were scaled using ScaleData, and module eigengenes were calculated using the ModuleEigengenes function and retrieved using the GetMEs function for downstream comparative analyses. Intramodular connectivity and hub gene identification were performed using the ModuleConnectivity and GetHubGenes functions, respectively, enabling characterization of module-specific transcriptional programs and highly connected genes within uveal melanocyte populations.

Functional enrichment analyses of module-associated genes were performed using Gene Set Enrichment Analysis (GSEA) and Gene Ontology (GO) enrichment analysis implemented through the GSEA and enrichGO functions from the clusterProfiler package (version 4.14.6) using the human (org.Hs.eg.db) annotation database. Multiple hypothesis testing correction was performed using false discovery rate (FDR) adjustment, and pathways with significance less than 0.05 were considered enriched. Differential module eigengene analyses between primary UM subgroups were performed using the FindDMEs function.

Comparisons were conducted according to tumor class status and *PRAME* expression status to identify modules associated with molecular risk stratification and metastatic potential in primary UM samples. Results were plotted with PlotDMEsLollipop function.

### Cellular communication network analysis

CellChat (version 2.1.2) was utilized to construct the cellular communication network from RNA counts of each scRNA-seq sample, excluding UMM072 and UMM075 due to limited cellular diversity. The CellChat human and mouse ligand-receptor interaction databases were used to construct communication networks. Average gene expression per group was determined by trimean, computing communication probabilities with cell population size considered requiring at least 10 cells per cellular group. Significant signaling in the generated cellular communication network was determined by comparing (total, sent, and received) signal probabilities by two-tailed Wilcoxon test via the stats R package. Differential signaling plots and differential ligand-receptor bubble plots were generated with the CellChat functions, netAnalysis_signalingChanges_scatter and netVisual_bubble, respectively.

### Single-cell TCR sequencing and analysis

Human filtered contigs were generated with CellRanger (version 5.0.0) with 10X Genomics GRCh38 human VDJ reference (version = “alts-ensembl-7.1.0”). Downstream quality assessment, repertoire characterization, and clonality analysis was conducted with scRepertoire (version 1.8.0). For each sample, TCR contigs were combined into clones with the combineTCR function from the scRepertoire package, filtering for multiple clones by selecting the 2 corresponding chains with the highest expression for a single cell barcode. The combineExpression function was used to merge TCR repertoire analysis data with scRNA-seq Seurat object and to generate categorical calls for clone size, setting the maximum TCR clone size bins for small, medium, large, and hyperexpanded up to 1%, 5%, 10%, and 100% of the sample repertoire, respectively. Strict nucleotide-level clone calls for both chains were used for clone calls. Clonal homeostasis, including assessment of clone size proportions, was visualized with the clonalHomeostasis function. Clonal diversity metrics were calculated and visualized with the clonalDiversity function. Comparison and visualization of shared versus unique clones and their relative abundances between samples was conducted with the clonalScatter function, using strict clonal calls.

### Spatial transcriptomic analysis

Fresh frozen OCT-embedded tumors were cryo-sectioned with Leica Biosystems Cryostat and placed within the 5×5mm capture areas of the 10X Visium Gene Expression (v1) slide. Tissue underwent hematoxylin and eosin (H&E) staining and imaged at 20X magnification with EVOS M7000 Imaging System (Thermo Fisher). Tissues were processed as recommended by the manufacturer, with 12 minutes of permeabilization. After sequencing, reads and images were initially processed with 10X Space Ranger (version 1.2.2). Capture dot quality control and plotting was conducted with Seurat. Spatial capture dot RNA expression was processed with the Load10X_Spatial function and transformed with the SCTransform function. PCA of spatial data was conducted on transformed expression data with RunPCA Seurat function. ElbowPlot Seurat function was used to identify significant principal components for each sample, with the first 7, 5, and 10 principal components used for UMM079, UMM085, and UMM086, respectively.

Clustering and UMAP-based dimension reduction was conducted with FindNeighbors, FindClusters, and RunUMAP Seurat functions, with clustering visualized at resolutions of 0.4 and 1. Log normalization and scaling of spatial expression data was conducted with NormalizeData and ScaleData Seurat functions. In addition to Seurat functions such as Dimplot and Dotplot, visualization was conducted with the FeatureScatter_scCustom and Plot_Density_Custom functions from the scCustomize R package.

Prediction of cell type annotation and transcriptional states for spatial capture dots was generated with FindTransferAnchors and TransferData Seurat functions using annotated tumor-matched scRNA-seq samples as reference with transformed reads. Cell type signature scores were generated with the AddModuleScore_UCell function from the UCell R package set to 1,500 maximum gene ranks. The evaluated cell type signatures and their associated marker genes included UM cells (*PMEL*, *MLANA*, *TYR*, *MITF*), fibroblasts (*PDGFRA*, *LUM*), smooth muscle cells (SMCs) (*MYH11*, *TAGLN*, *DES*), myofibroblasts (*PDGFRA*, *DCN*, *ACTA2*, *MYH11*), pericytes (*RGS5*, *CSPG4*, *MCAM*), monocytes (*CD14*, *FCGR3A*, *CCR2*), T cells (*CD3E*, *CD3G*, *CD27*, *CD28*, *SELL*), natural killer (NK) cells (*IL2RB*, *ZNF683*, *FGFBP2*, *KLRF1*), plasmacytoid dendritic cells (pDCs) (*PTPRC*, *IL3RA*, *CLEC4C*), CD8^+^ T cells (*CD8A*, *GZMA*, *GZMB*, *GZMK*), CD4^+^ T cells (*CD4*, *CD27*, *TCF7*), exhausted cells = c(*LAG3*, *PDCD1*, *TIGIT*, *HAVCR2*, *CTLA4*), regulatory T cells (Tregs) (*IL7R*, *FOXP3*, *IL2RA*, *CTLA4*), B cells (*CD19*, *CD79A*, *MS4A1*), plasma cells (*IGHG1*, *CD79A*, *MZB1*, *SDC1*), macrophages (*CD68*, *ITGAM*, *CD80*, *CD163*, *CD79*, *MSR1*), and endothelial cells (*VWF*, *PECAM1*, *CD34*). More detailed cell type annotations for spatial capture dots were called based on maximum cell type signature score(s), while the remainder without a maximum score were called as UM cells.

Capture dots with UM cell signature scores greater than 75^th^ percentile of all capture dots were labelled as UM cells. Capture dots annotated as UM cells were labelled as PRAME-positive if *PRAME* expression for that capture dot was positive. Broad cell type annotation of spatial capture dots was based on categorical grouping of detailed cell type annotations, where fibroblasts, pericytes, SMCs, and myofibroblasts were labelled as stroma, CD4^+^ T cells, CD8^+^ T cells, and Tregs were labelled as T cells, NK cell labels were retained, B cells, plasma cells, and pDCs were labelled as B lymphoid, monocytes and macrophages were labelled as myeloid, and endothelial cells were labelled as an endothelial-like category.

Giotto R package (version 1.0.4) was used to generate spatial proximity enrichment analysis. Processed spatial counts, spatial annotation, and tissue coordinates were used with the Giotto functions, createGiottoInstructions and createGiottoObject. The generated dataset was filtered with the filterGiotto function, requiring an expression threshold of 1, a minimum of 10 capture dots expressing a given gene, and a minimum of 10 genes detected in a capture dot. Filtered data were normalized with normalizeGiotto function with a scale factor of 1000. Spatial data was further processed for clustering with the Giotto functions, addStatistics, adjustGiottoMatrix, calculateHVG, and runPCA. Significant principal components were determined by jackstrawPlot function and used with the runUMAP Giotto function. Delaunay networks and shared nearest neighbors were generated with the createSpatialNetwork and createNearestNetwork Giotto functions. Delaunay network and transferred annotation was used for spatial interaction enrichment/depletion analysis of broad cell type interactions with cellProximityEnrichment Giotto function, set to calculate 10,000 simulations and adjust significance with FDR-correction. The Giotto functions, cellProximityBarplot and cellProximitySpatPlot2D, were utilized for visualizing results.

Spatially derived *PRAME* correlation analysis was conducted with all filtered spatial capture dots for UMM086, comparing log-normalized RNA expression of all genes to *PRAME* to calculate Spearman’s rank correlation with the cor.test function from the stats package. *P*-values were adjusted with Bonferroni correction. PRAME-associated genes were ranked by correlation coefficient and filtered for Bonferroni-adjusted *P*-values less than 0.05 and positive correlation with PRAME. A ranked list of genes associated with *PRAME* expression was analyzed by gene set enrichment analysis (GSEA) using the GSEA function in the clusterProfiler package, with the analysis restricted to positive enrichment scores (scoreType = “pos”) and human pathway gene sets obtained from MSigDB. *P* values in GSEA underwent FDR correction, and select pathways were visualized with the gseaplot2 function from the enrichplot R package.

### Cell culture

UM cell lines 92.1, Mel202 and Mel290 were maintained in RPMI medium supplemented with 10% tetracycline-free, heat-inactivated fetal bovine serum (FBS), 2 mM GlutaMAX and 1× penicillin–streptomycin in a humidified atmosphere containing 5% CO_2_. The non-transformed human uveal melanocyte cell line (UMC026) was generated from an enucleated human eye as previously described^35^ and were maintained in Ham’s F-12 medium under 5% CO_2_ and 5% O_2_ and supplemented with 10% tetracycline-free, heat-inactivated FBS, 2 mM GlutaMAX, 1× penicillin–streptomycin, 100 µM 3-isobutyl-1-methylxanthine (IBMX, PeproTech), basic fibroblast growth factor (bFGF; 10 ng ml^−1^; PeproTech), recombinant stem cell factor (rSCF; 10 ng ml^−1^; PeproTech) and epidermal growth factor (EGF; 20 ng ml^−1^; PeproTech). *BAP1*-knockout UMC026 cells were generated by CRISPR–Cas9-mediated biallelic deletion of exon 1 of *BAP1* in UMC026 cells^35^. 92.1 and Mel202 cells were engineered to express a doxycycline-inducible short hairpin RNA (shRNA) targeting *BAP1*, as previously described^35^. Mel290 and UMC026 cells were engineered for doxycycline-inducible *PRAME* expression, as previously described^36^. All cell lines were routinely tested for mycoplasma contamination and confirmed to be mycoplasma negative using a commercially available mycoplasma detection kit (ab289834, Abcam).

### Western blotting

BAP1-knockout cells and their corresponding wild-type controls were analyzed under basal culture conditions, whereas doxycycline-inducible *BAP1*-knockdown and *PRAME*-expressing cells were treated with doxycycline (1 µg ml^−1^; Thermo Fisher Scientific, J60579.14) for 72 h before protein extraction. Cells were washed with ice-cold phosphate-buffered saline (PBS) and lysed in RIPA buffer supplemented with protease and phosphatase inhibitors. Protein concentrations were determined using the Bradford assay, and 20–25 µg of total protein from each sample was resolved by SDS–PAGE and transferred to nitrocellulose membranes. Membranes were blocked in 5% bovine serum albumin (BSA; Thermo Fisher Scientific, J65731.10) in Tris-buffered saline containing 0.1% Tween-20 (TBS-T) and incubated overnight at 4 °C with primary antibodies against BAP1 (1:1,000; Santa Cruz Biotechnology, sc-28383), PRAME (1:1,000; Abcam, ab219650), β-actin (1:5,000; Invitrogen, MA1-140) or GAPDH (1:5,000; Santa Cruz Biotechnology, sc-47724). After washing with TBST, membranes were incubated for 1 hour at room temperature with the appropriate horseradish peroxidase-conjugated secondary antibodies (Bio-Rad, 1706515 and 1706516). Immunoreactive proteins were detected using Pierce ECL Plus Western Blotting Substrate (Thermo Fisher Scientific, 32132) and imaged using an iBright FL1500 Imaging System (Thermo Fisher Scientific).

### Quantitative reverse transcription PCR

BAP1-knockout cells and their corresponding wild-type controls were analyzed under basal culture conditions, whereas doxycycline-inducible *BAP1*-knockdown and *PRAME*-expressing cells were cultured with or without doxycycline (1 µg ml^−1^) for 72 hours before RNA extraction. Total RNA was extracted using TRIzol reagent (Invitrogen, 15596018), and 1 µg of RNA was used for cDNA synthesis with the Maxima First Strand cDNA Synthesis Kit (K1642, Thermo Fisher Scientific). Quantitative real-time PCR (qPCR) was performed using PowerUp SYBR Green Master Mix (Thermo Fisher Scientific, 3398048) on a QuantStudio 6 Pro Real-Time PCR System (Thermo Fisher Scientific). Relative mRNA expression was normalized to *18S rRNA*. Primer sequences are provided in Source Data.

## DATA AVAILABILITY

Data generated for analysis were made publicly accessible at Dryad Data Repository [https://doi.org/10.5061/dryad.7h44j109r], which includes access to Source Data.

## CODE AVAILABILITY

This study uses existing tools as described in the Methods sections.

## ACKNOWLEDGMENTS

This research was supported by Cancer Prevention and Research Institute of Texas Recruitment of Established Investigator Award RR220010 (J.W.H.); National Institute of General Medical Sciences Grant T32 GM145462 to the Medical Scientist Training Program of University of Miami (J.J.D.); NCI Cancer Center Support Grants P30 CA142543 to University of Texas Southwestern Simmons Comprehensive Cancer Center and P30 CA240139 to University of Miami Sylvester Comprehensive Cancer Center; NEI Center Core Grants P30 EY030413 to University of Texas Southwestern Department of Ophthalmology and P30 EY014801 to University of Miami Department of Ophthalmology; Research to Prevent Blindness, Inc. Challenge Grant to University of Texas Southwestern Department of Ophthalmology; and Research to Prevent Blindness, Inc. Unrestricted Grant to University of Miami Department of Ophthalmology.

## CONTRIBUTIONS

Conceptualization, J.W.H.; data collection, J.J.D., C.D., S.S., A.C., M.S., Z.M.C., J.W.H.; data analysis, J.J.D., M.S., X.L., J.H., J.W.H.; provision of resources, Z.M.C., J.W.H.; manuscript writing, J.J.D., J.W.H.; manuscript review and editing, all authors.

