## Supplementary Information for "BAP1 loss and *PRAME* expression converge to remodel the tumor–immune ecosystem during uveal melanoma progression"

**SUPPLEMENTARY FIGURES**

**Supplementary Fig. 1 | Chromosome-scale copy-number profiles of Class 1 and Class 2 uveal melanoma cells.** Heat maps showing chromosome-scale copy-number profiles inferred from single-cell RNA sequencing data using inferCNV. Uveal melanoma (UM) cells from Class 1 and Class 2 primary tumors were analyzed separately using identical inferCNV parameters and a shared reference population of 2,500 adult human choroidal melanocytes. Rows represent individual cells and columns represent genes ordered by genomic position and grouped by chromosome. The upper heat map shows the shared melanocyte reference population, whereas the middle and lower heat maps show Class 1 ($n$ = 11,236) and Class 2 ($n$ = 14,167) UM cells, respectively. Red and blue indicate relative increases and decreases in chromosome-smoothed gene expression compared with the reference population, consistent with inferred copy-number gains and losses, respectively; white indicates an approximately neutral signal. Vertical black lines demarcate chromosomes. Dendrograms show hierarchical clustering of UM cells and adjacent annotation strips denote inferCNV-defined subclusters and single-cell RNA sequencing-defined tumor transcriptional states.

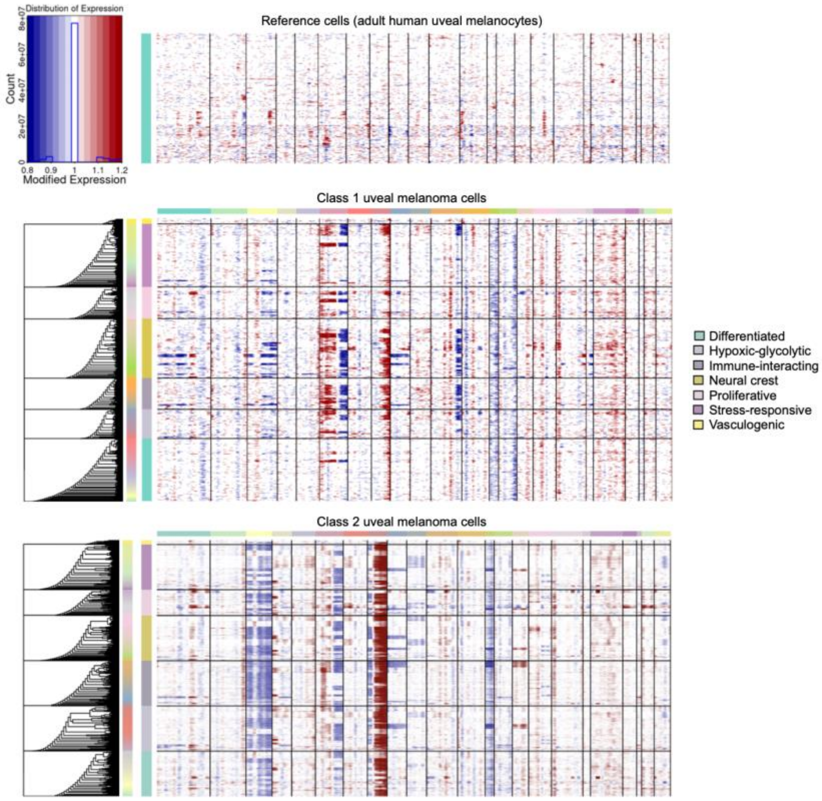

**Supplementary Fig. 2 | Western blot validation of BAP1 depletion and *PRAME* induction in uveal melanocytic cell lines. a**, Western blot analysis of BAP1 protein in the non-malignant uveal melanocytic cell line UMC026 with wild-type BAP1 (WT) or engineered BAP1 knockout (KO). **b**, Western blot analysis of BAP1 protein in the 92.1 and Mel202 uveal melanoma (UM) cell lines engineered with doxycycline-inducible BAP1 knockdown (KD) and cultured without (−) or with (+) doxycycline (Dox). **c**, Western blot analysis of PRAME protein in UMC026 and Mel290 cells engineered with doxycycline-inducible PRAME expression and cultured without (−) or with (+) Dox. β-actin served as the loading control in **a,b**, and GAPDH served as the loading control in **c**. Cropped blots are shown above their corresponding uncropped images. Red boxes indicate the regions displayed in the cropped blots. Molecular masses are indicated in kilodaltons (kDa).

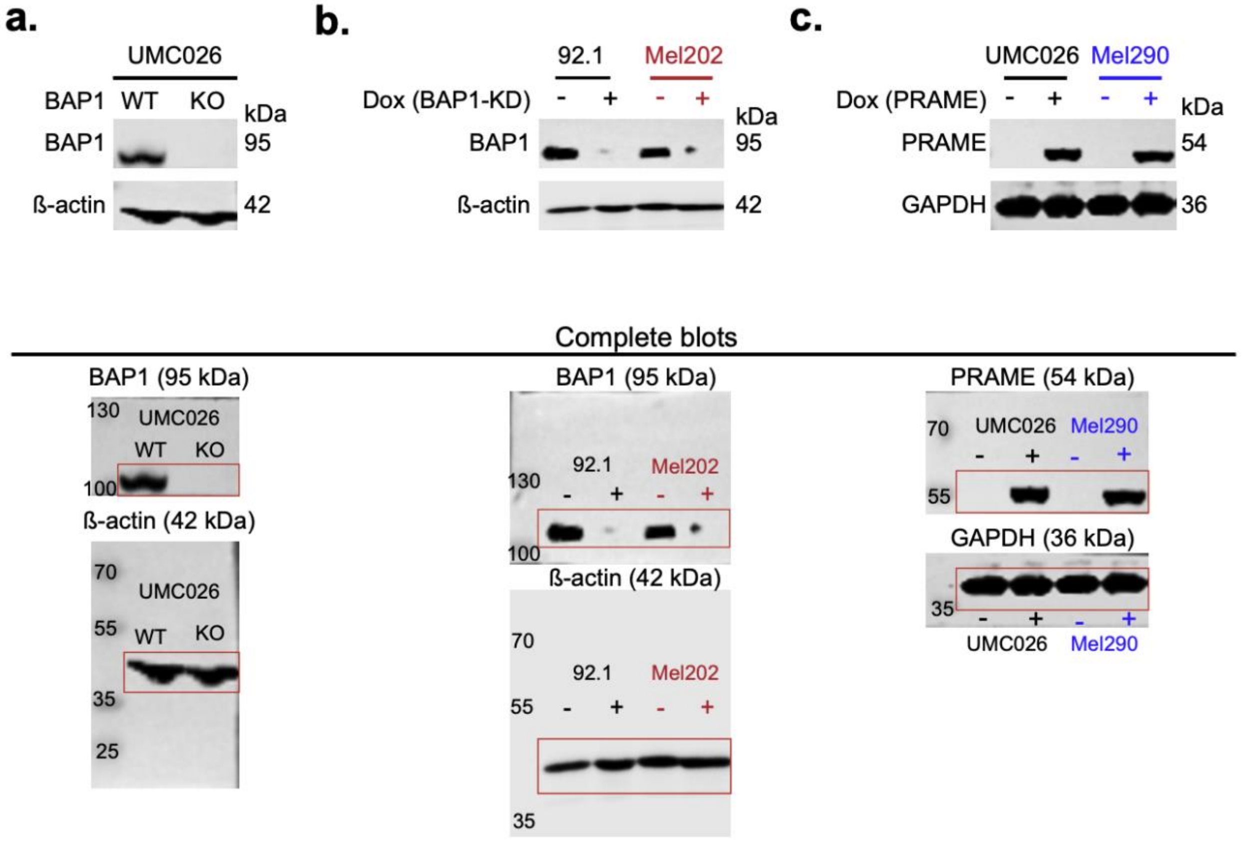

**Supplementary Fig. 3 | Distribution of M1-like and M2-like macrophages across uveal melanoma molecular subgroups.** Violin plots showing the proportions of tumor-associated macrophages (TAMs) classified as M1-like, M2-like or M1/M2-negative. The analysis included 4,174 TAMs from the single-cell RNA sequencing cohort, including 32 tumor samples from 25 independent tumors grouped by molecular subtype based on 15-gene expression profiling (15-GEP) and *PRAME* expression status for primary tumors: Class 1/PRAME(−) (*n* = 3), Class 1/PRAME(+) (*n* = 4), Class 2/PRAME(−) (*n* = 3) and Class 2/PRAME(+) (*n* = 7). TAMs from 8 metastatic samples are shown separately. UCell signature scores were calculated for each macrophage using a maximum rank of 1,500 genes. The M1 signature comprised *CD80*, *TNF*, *IL1A* and *CD86*, whereas the M2 signature comprised *CD163*, *MRC1* and *CD209*. TAMs were classified as M2-like when the M2 signature score was positive and greater than or equal to the M1 signature score, or as M1-like when the M1 signature score was positive and greater than the M2 signature score. TAMs with scores of zero for both signatures were classified as M1/M2-negative. For each sample, abundance was calculated as the proportion of all TAMs assigned to the indicated category. Violin width represents the estimated density of sample-level proportions. Boxes show the median and interquartile range (IQR), whiskers extend to 1.5 × the IQR, and each point represents one sample. Statistical significance was assessed using two-sided Wilcoxon rank-sum tests.

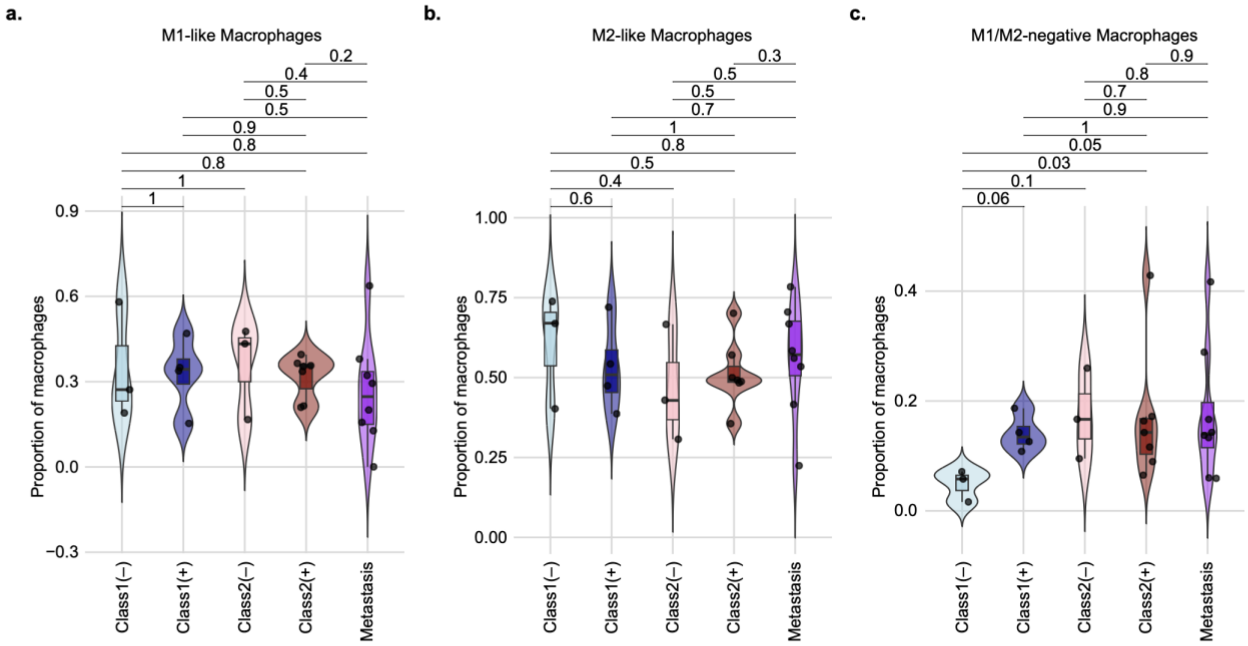

### SUPPLEMENTARY TABLES

**Supplementary Table 1 | Molecular and sequencing characteristics of uveal melanoma and reference samples analyzed in this study.** The GEP Class and PRAME columns indicate 15-gene expression profile (15-GEP) class (Class 1 or Class 2) and *PRAME* expression status (negative or positive), respectively. GEP class reported for metastatic samples refers to molecular profiling of the corresponding primary tumor. The Gq/11 mutation column reports the initiating *GNAQ* or *GNA11* driver mutation and the corresponding amino acid substitution when available (Q209L, Q209P or R183C). The BSE mutation column reports secondary driver mutations involving *BAP1*, *SF3B1* or *EIF1AX*. The Library Capture column specifies whether 3′ or 5′ gene expression capture chemistry was used. The Sample Source column indicates whether each dataset was newly generated in this study or obtained from a previously published source. Abbreviations: N/A, not applicable.

| Sample ID | Sample Descriptions | GEP Class | PRAME | Gq/11 Mutation | BSE Mutation | Library Capture Orientation | Sample Source |
|---|---|---|---|---|---|---|---|
| UMM059 | UMM059 Primary | 2 | Positive | GNA11 (Q209P) | BAP1 (V616fs) | 3' | Durante et al 2020 |
| UMM061 | UMM061 Primary | 2 | Positive | GNA11 (Q209L) | BAP1 (S123fs) | 5' | Durante et al 2020 |
| UMM063 | UMM063 Primary | 2 | Negative | GNAQ (Q209L) | BAP1 (null) | 5' | Durante et al 2020 |
| UMM064 | UMM064 Primary | 2 | Positive | GNAQ (Q209P) | BAP1 (S430fs) | 5' | Durante et al 2020 |
| UMM065 | UMM065 Primary | 1 | Negative | GNA11 (Q209L) | EIF1AX (N4Y) | 5' | Durante et al 2020 |
| UMM066 | UMM066 Primary | 2 | Positive | GNAQ (Q209P) | BAP1 (E125spice) | 5' | Durante et al 2020 |
| UMM069 | UMM069 Primary | 2 | Positive | GNAQ (Q209L) | BAP1 (R146splice) | 5' | Durante et al 2020 |
| UMM071 | UMM071 Primary | 2 | Negative | GNA11 (R183C) | BAP1 | 5' | New case |
| UMM072 | UMM072 Primary | 2 | Negative | GNAQ (Q209P) | BAP1 (V209fs) | 5' | New case |
| UMM075 | UMM075 Primary | 2 | Negative | GNA11 (R183C) | BAP1 (R179W) | 5' | New case |
| UMM076 | UMM076 Primary | 1 | Positive | GNAQ (Q209P) | SF3B1 (R625H) | 5' | New case |
| UMM079A | UMM079 Multiregional (anterior) primary | 1 | Negative | GNAQ (Q209P) | EIF1AX | 5' | New case |
| UMM079M | UMM079 Multiregional (middle) primary | 1 | Negative | GNAQ (Q209P) | EIF1AX | 5' | New case |
| UMM079P | UMM079 Multiregional (posterior) primary | 1 | Negative | GNAQ (Q209P) | EIF1AX | 5' | New case |
| UMM080intra | UMM080 Intraocular primary | 2 | Positive | GNA11 (Q209L) | Unknown | 5' | New case |
| UMM080extra | UMM080 Extraocular extension of primary | 2 | Positive | GNA11 (Q209L) | Unknown | 5' | New case |
| UMM081 | UMM081 Primary | 1 | Negative | GNA11 (Q209L) | BAP1/EIF1AX | 5' | New case |
| UMM084apex | UMM084 Multiregional (apex) primary | 2 | Positive | GNAQ (Q209L) | BAP1 | 5' | New case |
| UMM084base | UMM084 Multiregional (base) primary | 2 | Positive | GNAQ (Q209L) | BAP1 | 5' | New case |
| UMM085apex | UMM085 Multiregional (apex) primary | 1 | Positive | GNA11 (Q209L) | SF3B1 | 5' | New case |
| UMM085base | UMM085 Multiregional (base) primary | 1 | Positive | GNA11 (Q209L) | SF3B1 | 5' | New case |
| UMM086ant | UMM086 Multiregional (anterior) primary | 1 | Positive | GNAQ (Q209P) | SF3B1/EIF1AX | 5' | New case |
| UMM086post | UMM086 Multiregional (posterior) primary | 1 | Positive | GNAQ (Q209P) | SF3B1/EIF1AX | 5' | New case |
| UMM088lat | UMM088 Multiregional (lateral) primary | 1 | Positive | GNAQ (Q209P) | SF3B1 | 5' | New case |
| UMM088med | UMM088 Multiregional (medial) primary | 1 | Positive | GNAQ (Q209P) | SF3B1 | 5' | New case |
| BSSR0022 | BSSR0022 Liver metastasis | 1 | Positive | GNAQ (Q209P) | SF3B1 (R625H) | 5' | Durante et al 2020 |
| UMM067L | UMM067 Liver metastasis | 2 | Positive | GNA11 (Q209L) | BAP1 (G220splice) | 5' | Durante et al 2020 |
| UMM041L | UMM041 Liver metastasis | 2 | Positive | GNAQ (Q209P) | BAP1 (D311fs) | 5' | Durante et al 2020 |
| UMM072L | UMM072 Liver metastasis | 2 | Negative | GNAQ (Q209P) | BAP1 (V209fs) | 5' | New case |
| FNA0194L | FNA0194 Liver metastasis | 2 | Positive | Unknown | Unknown | 5' | New case |
| UMM005L | UMM005 Liver metastasis | 2 | Positive | Unknown | Unknown | 5' | New case |
| FNA0257SM | FNA0257 Subcutaneous metastasis | 1 | Positive | Unknown | BAP1 (null) | 5' | New case |
| UMM072SM | UMM072 Subcutaneous metastasis | 2 | Negative | GNAQ (Q209P) | BAP1 (V209fs) | 5' | New case |
| UMC071M | UMC071 Congenital choroidal nevus | N/A | N/A | GNA11 (R183C) | N/A | 5' | New case |
| adult1 | Adult choroid 1 replicate 1 | N/A | N/A | N/A | N/A | 3' | Collin et al 2023 |
| adult1rep2 | Adult choroid 1 replicate 2 | N/A | N/A | N/A | N/A | 3' | Collin et al 2023 |
| adult1rep3 | Adult choroid 1 replicate 3 | N/A | N/A | N/A | N/A | 3' | Collin et al 2023 |
| adult1rep4 | Adult choroid 1 replicate 4 | N/A | N/A | N/A | N/A | 3' | Collin et al 2023 |
| adult2 | Adult choroid 2 replicate 1 | N/A | N/A | N/A | N/A | 3' | Collin et al 2023 |
| adult2rep2 | Adult choroid 2 replicate 2 | N/A | N/A | N/A | N/A | 3' | Collin et al 2023 |
| adult2rep3 | Adult choroid 2 replicate 3 | N/A | N/A | N/A | N/A | 3' | Collin et al 2023 |
| adult2rep4 | Adult choroid 2 replicate 4 | N/A | N/A | N/A | N/A | 3' | Collin et al 2023 |
| adult3 | Adult choroid 3 replicate 1 | N/A | N/A | N/A | N/A | 3' | Collin et al 2023 |
| adult3rep2 | Adult choroid 3 replicate 2 | N/A | N/A | N/A | N/A | 3' | Collin et al 2023 |
| adult4 | Adult choroid 4 | N/A | N/A | N/A | N/A | 3' | Collin et al 2023 |
| adult5 | Adult choroid 5 replicate 1 | N/A | N/A | N/A | N/A | 3' | Collin et al 2023 |
| adult5rep2 | Adult choroid 5 replicate 2 | N/A | N/A | N/A | N/A | 3' | Collin et al 2023 |
| fetal12PCW | Fetal choroid 12 Post-conception Weeks (PCW) replicate 1 | N/A | N/A | N/A | N/A | 3' | Collin et al 2023 |
| fetal12PCWrep2 | Fetal choroid 12 PCW replicate 2 | N/A | N/A | N/A | N/A | 3' | Collin et al 2023 |
| fetal16PCW | Fetal choroid 16 PCW replicate 1 | N/A | N/A | N/A | N/A | 3' | Collin et al 2023 |
| fetal16PCWrep2 | Fetal choroid 16 PCW replicate 2 | N/A | N/A | N/A | N/A | 3' | Collin et al 2023 |
| fetal20PCW | Fetal choroid 20 PCW replicate 1 | N/A | N/A | N/A | N/A | 3' | Collin et al 2023 |
| fetal20PCWrep2 | Fetal choroid 20 PCW replicate 2 | N/A | N/A | N/A | N/A | 3' | Collin et al 2023 |
| fetal21PCW | Fetal choroid 21 PCW replicate 1 | N/A | N/A | N/A | N/A | 3' | Collin et al 2023 |
| fetal21PCWrep2 | Fetal choroid 21 PCW replicate 2 | N/A | N/A | N/A | N/A | 3' | Collin et al 2023 |

**Supplementary Table 2 | Positive and negative marker gene signatures used for UCell-based cell type annotation.** The first column identifies the cell type represented by each signature. Marker genes are listed by gene symbol, followed by a plus sign (+) for positive markers or a minus sign (−) for negative markers. The Samples with Signature Scored column specifies the samples in which each signature was applied during cell type annotation: all samples, ocular samples only, liver metastasis samples only or all metastatic samples.

| UCell Cell Type Signature ID | UCell Cell Type Signature Description | Positive and Negative Markers | Samples with Signature Scored |
|---|---|---|---|
| Melanocytic_sig | Melanocytic cell type signature | "MLANA+","PMEL+","TYRP1+", "TYR+", "PAX3+" | All |
| Tcell_sig | T cell signature | "PMEL-","MLANA-","CD2+","CD3E+","CD3D+" | All |
| CD8Tcell_sig | CD8 T cell signature | "PMEL-","MLANA-","CD2+","CD3E+","CD8A+","CD8B+","CD4-" | All |
| CD4Tcell_sig | CD4 T cell signature | "PMEL-","MLANA-","CD2+","CD3E+","CD4+","CD8-" | All |
| Treg_sig | Regulatory T cell (Treg) signature | PMEL-,"MLANA-","CD3E+", "CD3D+", "CD4+", "FOXP3+", "CTLA4+" | All |
| Bcells_sig | B cell signature | "PMEL-","MLANA-","CD19+","CD79A+","MS4A1+" | All |
| Plasma_sig | Plasma cell signature | "PMEL-","MLANA-","IGHG1+","MZB1+","SDC1+","CD79A+","JCHAIN+" | All |
| Macrophage_sig | Macrophage signature | "PMEL-","MLANA-","CD68+","FCGR1A+","TYROBP+" | All |
| Monocyte_sig | Monocyte signature | "PMEL-","MLANA-","CD14+","FCGR1A-","CD68-" | All |
| NKcell_sig | NK cell signature | "PMEL-","MLANA-","CD3E-","IL2RB+","ZNF683+","FGFBP2+","SPON2+","KLRF1+","FCGR3A+","TYROBP+" | All |
| Endothelial_sig | Endothelial cell signature | "PMEL-","MLANA-","VWF+","PECAM1+","PMEL-","MLANA-" | All |
| HSCs_sig | Hepatic stellate cell (HSC) signature | "PMEL-","MLANA-","CD3E-","TIMP1+","VCL+","IGFBP6+" | Liver metastasis |
| Fibroblast_sig | Fibroblast signature | "PMEL-","MLANA-","CD3E-","PMEL-","MLANA-","PDGFRA+","LUM+","MYH11-" | All |
| MyoFibro_sig | Myofibroblast signature | "PMEL-","MLANA-","CD3E-","PMEL-","MLANA-","PDGFRA+","DCN+","ACTA2+","MYH11+","DES+","TAGLN+" | All |
| Pericytes_sig | Pericytes signature | "PMEL-","MLANA-","CD3E-","PMEL-","MLANA-","PDGFRA+","RGS5+" | All |
| MullerCells_sig | Muller cell signature | "PMEL-,"MLANA-", "TYROBP-", "OAF+", "GLUL+", "DKK3+" | Ocular |
| SMCs_sig | Smooth muscle cell (SMC) signature | "PMEL-","MLANA-","PDGFRA-","LUM-","MYH11+","TAGLN+","SH3BGR+","DES+","SMTN+" | All |
| Photorecept_sig | Photoreceptor signature | "PMEL-","MLANA-","RCVRN+","CRX+" | Ocular |
| RPE_sig | Retinal pigment epithelium (RPE) signature | "RPE65+","RLBP1+","BEST1+" | Ocular |
| Hepatocyte_sig | Hepatocyte signature | "PMEL-","MLANA-","CYP2E1+","KRT8+","KRT18+","ALB+" | Liver metastasis |
| Erythoid_sig | Erythroid-like cell signature | "PMEL-","MLANA-","HBB+","HBA1+","HBA2+" | All |
| pDC_sig | Plasmacytoid dendritic cell (pDC) signature | "PMEL-","MLANA-","PTPRC+","IL3RA+","CLEC4C+","JCHAIN+" | All |
| cDC_sig | Conventional dentritic cell (cDC) signature | "PMEL-","MLANA-","ZBTB46+","LAMP3+","ITGAX+","CD33+" | All |
| Mastcells_sig | Mast cell signature | "PMEL-","MLANA-","CD4-","CD8A-","KIT+","TPSAB1+","IL1RL1+" | All |
| Schwann_sig | Schwann cell signature | "PMEL-","MLANA-","MPZ+","PLP1+" | Metastasis |
| RetinalGanglionCells_sig | Retinal ganglion cell (RGC) signature | "PMEL-","MLANA-", "ISL1+", "STMN2+" | Ocular |
| Astrocytes_sig | Astrocytes signature | "PMEL-","MLANA-", "NGFR+", "GFAP+" | Ocular |

**Supplementary Table 3 | Curated gene signatures used for UCell-based annotation of transcriptional states in uveal melanoma cells.** The table lists the marker genes comprising signatures for the seven uveal melanoma (UM) cell transcriptional states: differentiated, stress-responsive, neural crest-like, hypoxic–glycolytic, immune-interacting, proliferative and vasculogenic. The table also includes metrics for ranking cluster-associated differentially expressed genes (DEGs) and a description of the marker selection basis. Additional details are provided in Methods. These signatures were used to calculate UCell scores for transcriptional-state annotation of individual UM cells. Genes are listed by gene symbol.

Abbreviations: pct.1, percent expression in group 1 (cluster of interest); pct.2, percent expression in group 2 (all other clusters); log2FC, log2 fold change.

| Transcriptional state | Gene | Supporting cluster | P value | Average log2FC | pct.1 | pct.2 | pct.1 - pct.2 | Bonferroni-adjusted P | Selection status (Exception criterion) |
|---|---|---|---|---|---|---|---|---|---|
| Differentiated | CD82 | 0 | 0 | 0.879682041 | 0.584 | 0.332 | 0.252 | 0 | Top-20 marker |
| Differentiated | ENPP2 | 0 | 0 | 0.841649679 | 0.417 | 0.229 | 0.188 | 0 | Top-20 marker |
| Differentiated | LTA4H | 0 | 6.4942E-286 | 0.514592754 | 0.515 | 0.378 | 0.137 | 1.9667E-281 | Curated exception (pct.1 - pct.2 < 0.15) |
| Differentiated | AZGP1 | 8 | 9.2638E-196 | 0.75175426 | 0.422 | 0.243 | 0.179 | 2.8055E-191 | Top-20 marker |
| Differentiated | GNG11 | 12 | 0 | 1.215213532 | 0.908 | 0.61 | 0.298 | 0 | Top-20 marker |
| Differentiated | LNP1 | 0 | 0 | 0.638574807 | 0.664 | 0.446 | 0.218 | 0 | Top-20 marker |
| Differentiated | RAB32 | 0 | 0 | 0.624084372 | 0.739 | 0.573 | 0.166 | 0 | Top-20 marker |
| Differentiated | HHATL | 0 | 0 | 0.908314786 | 0.412 | 0.218 | 0.194 | 0 | Top-20 marker |
| Differentiated | PPP1R3C | 0 | 0 | 0.824683565 | 0.382 | 0.209 | 0.173 | 0 | Top-20 marker |
| Differentiated | GALNTL6 | 0 | 0 | 0.991811664 | 0.289 | 0.13 | 0.159 | 0 | Top-20 marker |
| Differentiated | GPR37 | 12 | 0 | 1.052116055 | 0.68 | 0.275 | 0.405 | 0 | Top-20 marker |
| Differentiated | SSPN | 0 | 0 | 0.816582474 | 0.447 | 0.252 | 0.195 | 0 | Top-20 marker |
| Differentiated | ABRACL | 0 | 0 | 0.689176293 | 0.488 | 0.323 | 0.165 | 0 | Top-20 marker |
| Differentiated | ROPN1B | 0 | 0 | 0.578826576 | 0.441 | 0.267 | 0.174 | 0 | Top-20 marker |
| Stress-responsive | HSPA5 | 2 | 0 | 0.431511525 | 0.824 | 0.636 | 0.188 | 0 | Curated exception (average log2FC < 0.50) |
| Stress-responsive | ATF3 | 2 | 0 | 1.49700126 | 0.856 | 0.417 | 0.439 | 0 | Top-20 marker |
| Stress-responsive | DDIT3 | 2 | 0 | 1.492782455 | 0.82 | 0.42 | 0.4 | 0 | Top-20 marker |
| Stress-responsive | PPP1R15A | 2 | 0 | 1.519519332 | 0.961 | 0.6 | 0.361 | 0 | Top-20 marker |
| Stress-responsive | HERPUD1 | 2 | 0 | 0.806457176 | 0.755 | 0.524 | 0.231 | 0 | Passed filter; rank >20 |
| Stress-responsive | XBP1 | 2 | 0 | 0.667391076 | 0.639 | 0.427 | 0.212 | 0 | Passed filter; rank >20 |
| Stress-responsive | GADD45B | 2 | 0 | 1.5697989 | 0.748 | 0.391 | 0.357 | 0 | Top-20 marker |
| Stress-responsive | DNAJB1 | 2 | 0 | 1.686941987 | 0.946 | 0.607 | 0.339 | 0 | Top-20 marker |
| Stress-responsive | FOSB | 2 | 0 | 1.694362497 | 0.944 | 0.54 | 0.404 | 0 | Top-20 marker |
| Stress-responsive | ZFP36 | 2 | 0 | 1.60067845 | 0.946 | 0.552 | 0.394 | 0 | Top-20 marker |
| Stress-responsive | JUN | 5 | 0 | 1.254637494 | 0.971 | 0.758 | 0.213 | 0 | Top-20 marker |
| Stress-responsive | JUNB | 5 | 0 | 1.097609035 | 0.989 | 0.762 | 0.227 | 0 | Top-20 marker |
| Stress-responsive | EGR1 | 2 | 0 | 1.349261299 | 0.97 | 0.652 | 0.318 | 0 | Top-20 marker |
| Neural crest-like | HTR2B | 3 | 0 | 2.132608471 | 0.667 | 0.17 | 0.497 | 0 | Top-20 marker |
| Neural crest-like | PTP4A3 | 20 | 4.7428E-292 | 2.649507765 | 0.993 | 0.374 | 0.619 | 1.4363E-287 | Top-20 marker |
| Neural crest-like | IGFBP2 | 17 | 0 | 3.160740343 | 0.765 | 0.145 | 0.62 | 0 | Top-20 marker |
| Neural crest-like | FABP5 | 3 | 0 | 1.461359302 | 0.809 | 0.384 | 0.425 | 0 | Top-20 marker |
| Neural crest-like | TNFRSF12A | 17 | 0 | 3.180643078 | 0.776 | 0.149 | 0.627 | 0 | Top-20 marker |
| Neural crest-like | TRIB1 | 20 | 0 | 3.205376705 | 0.894 | 0.195 | 0.699 | 0 | Top-20 marker |
| Neural crest-like | RIMS2 | 22 | 0 | 3.493281938 | 0.677 | 0.048 | 0.629 | 0 | Top-20 marker |
| Neural crest-like | PROS1 | 22 | 2.8128E-275 | 4.146942763 | 0.898 | 0.165 | 0.733 | 8.5184E-271 | Top-20 marker |
| Neural crest-like | S100A4 | 15 | 0 | 3.669804669 | 0.807 | 0.111 | 0.696 | 0 | Top-20 marker |
| Neural crest-like | MIA | 15 | 0 | 2.641789134 | 0.852 | 0.157 | 0.695 | 0 | Top-20 marker |
| Neural crest-like | ADGRG1 | 15 | 0 | 1.732806535 | 0.883 | 0.301 | 0.582 | 0 | Top-20 marker |
| Neural crest-like | AHNAK2 | 22 | 6.4895E-120 | 1.349170787 | 0.764 | 0.17 | 0.594 | 1.9653E-115 | Passed filter; rank >20 |
| Neural crest-like | SPON2 | 17 | 0 | 2.452575825 | 0.933 | 0.393 | 0.54 | 0 | Passed filter; rank >20 |
| Hypoxic-glycolytic | SLC2A3 | 9 | 0 | 2.950567215 | 0.506 | 0.129 | 0.377 | 0 | Top-20 marker |
| Hypoxic-glycolytic | SLC16A3 | 9 | 0 | 3.458387185 | 0.352 | 0.059 | 0.293 | 0 | Top-20 marker |
| Hypoxic-glycolytic | PFKP | 9 | 0 | 1.873841111 | 0.327 | 0.135 | 0.192 | 0 | Passed filter; rank >20 |
| Hypoxic-glycolytic | PDK1 | 9 | 0 | 2.818120143 | 0.14 | 0.025 | 0.115 | 0 | Curated exception (pct.1 < 0.25; pct.1 - pct.2 < 0.15) |
| Hypoxic-glycolytic | AK4 | 9 | 0 | 2.226123014 | 0.285 | 0.093 | 0.192 | 0 | Passed filter; rank >20 |
| Hypoxic-glycolytic | NDUFA4L2 | 9 | 0 | 4.540580994 | 0.191 | 0.011 | 0.18 | 0 | Curated exception (pct.1 < 0.25) |
| Hypoxic-glycolytic | TMEM45A | 9 | 0 | 2.265185175 | 0.305 | 0.118 | 0.187 | 0 | Passed filter; rank >20 |
| Hypoxic-glycolytic | HSPB7 | 9 | 0 | 3.334816417 | 0.427 | 0.107 | 0.32 | 0 | Top-20 marker |
| Hypoxic-glycolytic | ERO1A | 9 | 0 | 2.525684732 | 0.3 | 0.118 | 0.182 | 0 | Passed filter; rank >20 |
| Hypoxic-glycolytic | SPAG4 | 9 | 0 | 3.960030777 | 0.186 | 0.02 | 0.166 | 0 | Curated exception (pct.1 < 0.25) |
| Hypoxic-glycolytic | NOL3 | 9 | 0 | 1.767962056 | 0.451 | 0.25 | 0.201 | 0 | Passed filter; rank >20 |
| Hypoxic-glycolytic | ADSSL1 | 9 | 9.4707E-254 | 2.073479746 | 0.336 | 0.175 | 0.161 | 2.8681E-249 | Passed filter; rank >20 |
| Immune-interacting | ISG15 | 7 | 0 | 3.845729386 | 0.589 | 0.119 | 0.47 | 0 | Top-20 marker |
| Immune-interacting | MX1 | 7 | 0 | 3.781968985 | 0.55 | 0.107 | 0.443 | 0 | Top-20 marker |
| Immune-interacting | STAT1 | 7 | 0 | 2.931031438 | 0.848 | 0.245 | 0.603 | 0 | Top-20 marker |
| Immune-interacting | GBP1 | 7 | 0 | 4.260711083 | 0.456 | 0.04 | 0.416 | 0 | Top-20 marker |
| Immune-interacting | IFIT3 | 7 | 0 | 3.33757891 | 0.528 | 0.131 | 0.397 | 0 | Passed filter; rank >20 |
| Immune-interacting | TAP1 | 7 | 0 | 3.130467782 | 0.83 | 0.211 | 0.619 | 0 | Top-20 marker |
| Immune-interacting | PSMB9 | 7 | 0 | 3.071655205 | 0.792 | 0.186 | 0.606 | 0 | Top-20 marker |
| Immune-interacting | TAP2 | 7 | 0 | 2.468497725 | 0.511 | 0.156 | 0.355 | 0 | Passed filter; rank >20 |
| Immune-interacting | HLA-DRA | 7 | 0 | 5.106416873 | 0.462 | 0.038 | 0.424 | 0 | Top-20 marker |
| Immune-interacting | HLA-DPB1 | 7 | 0 | 2.659260984 | 0.465 | 0.184 | 0.281 | 0 | Passed filter; rank >20 |
| Immune-interacting | HLA-DRB1 | 7 | 0 | 5.192224001 | 0.389 | 0.026 | 0.363 | 0 | Top-20 marker |
| Immune-interacting | HLA-DPA1 | 7 | 0 | 4.924305732 | 0.325 | 0.033 | 0.292 | 0 | Top-20 marker |
| Proliferative | UBE2C | 13 | 0 | 9.545651223 | 0.476 | 0.003 | 0.473 | 0 | Top-20 marker |
| Proliferative | PCLAF | 13 | 0 | 7.514560447 | 0.627 | 0.011 | 0.616 | 0 | Top-20 marker |
| Proliferative | RRM2 | 13 | 0 | 9.064187918 | 0.362 | 0.001 | 0.361 | 0 | Top-20 marker |
| Proliferative | CDK1 | 13 | 0 | 8.664094794 | 0.402 | 0.003 | 0.399 | 0 | Top-20 marker |
| Proliferative | PKMYT1 | 13 | 0 | 8.354254714 | 0.432 | 0.002 | 0.43 | 0 | Top-20 marker |
| Proliferative | BIRC5 | 13 | 0 | 8.317652646 | 0.436 | 0.004 | 0.432 | 0 | Top-20 marker |
| Proliferative | TYMS | 13 | 0 | 5.419682233 | 0.771 | 0.077 | 0.694 | 0 | Top-20 marker |
| Proliferative | TK1 | 13 | 0 | 7.804401809 | 0.461 | 0.004 | 0.457 | 0 | Top-20 marker |
| Proliferative | NUSAP1 | 13 | 0 | 7.852337594 | 0.447 | 0.008 | 0.439 | 0 | Top-20 marker |
| Proliferative | TOP2A | 13 | 0 | 8.306850669 | 0.357 | 0.003 | 0.354 | 0 | Top-20 marker |
| Proliferative | PBK | 13 | 0 | 8.534681123 | 0.355 | 0.001 | 0.354 | 0 | Top-20 marker |
| Proliferative | MKI67 | 13 | 0 | 8.703355886 | 0.327 | 0.002 | 0.325 | 0 | Top-20 marker |
| Proliferative | SPC25 | 13 | 0 | 8.505694581 | 0.268 | 0.001 | 0.267 | 0 | Top-20 marker |
| Proliferative | CCNA2 | 13 | 0 | 8.302864902 | 0.31 | 0.002 | 0.308 | 0 | Top-20 marker |
| Proliferative | CDC20 | 13 | 0 | 7.695618604 | 0.294 | 0.004 | 0.29 | 0 | Top-20 marker |
| Vasculogenic | COL1A1 | 19 | 0 | 11.99401366 | 0.67 | 0.002 | 0.668 | 0 | Top-20 marker |
| Vasculogenic | COL3A1 | 19 | 0 | 11.29055761 | 0.709 | 0.002 | 0.707 | 0 | Top-20 marker |
| Vasculogenic | MGP | 19 | 0 | 10.54843313 | 0.774 | 0.007 | 0.767 | 0 | Top-20 marker |
| Vasculogenic | DCN | 19 | 0 | 12.52447123 | 0.541 | 0.001 | 0.54 | 0 | Top-20 marker |
| Vasculogenic | OLFML3 | 19 | 0 | 10.39357099 | 0.602 | 0.003 | 0.599 | 0 | Top-20 marker |
| Vasculogenic | COL6A3 | 19 | 0 | 9.978680359 | 0.565 | 0.004 | 0.561 | 0 | Top-20 marker |
| Vasculogenic | FN1 | 19 | 0 | 7.987503559 | 0.774 | 0.034 | 0.74 | 0 | Top-20 marker |
| Vasculogenic | PDGFRB | 19 | 0 | 9.499274154 | 0.526 | 0.002 | 0.524 | 0 | Top-20 marker |
| Vasculogenic | RGS5 | 19 | 0 | 8.855781075 | 0.315 | 0.004 | 0.311 | 0 | Passed filter; rank >20 |
| Vasculogenic | TAGLN | 19 | 0 | 9.467093553 | 0.617 | 0.008 | 0.609 | 0 | Top-20 marker |
| Vasculogenic | ACTA2 | 19 | 4.0077E-127 | 6.844575567 | 0.348 | 0.072 | 0.276 | 1.2137E-122 | Passed filter; rank >20 |
| Vasculogenic | PLVAP | 21 | 0 | 12.90816172 | 0.445 | 0 | 0.445 | 0 | Top-20 marker |
| Vasculogenic | PECAM1 | 21 | 0 | 11.88669239 | 0.542 | 0.001 | 0.541 | 0 | Top-20 marker |
| Vasculogenic | CDH5 | 21 | 0 | 12.81771713 | 0.375 | 0 | 0.375 | 0 | Top-20 marker |
| Vasculogenic | ESAM | 21 | 0 | 10.03967813 | 0.441 | 0.001 | 0.44 | 0 | Top-20 marker |
